# Targeted sequencing and iterative assembly of near-complete genomes

**DOI:** 10.1101/2025.03.31.646505

**Authors:** Hasindu Gamaarachchi, Igor Stevanovski, Jillian M. Hammond, Andre L. M. Reis, Melissa Rapadas, Kavindu Jayasooriya, Tonia Russell, Dennis Yeow, Yvonne Hort, Chirag Patel, Andrew J. Mallett, Elaine Stackpoole, Lauren Roman, Luke W. Silver, Carolyn J. Hogg, Louise M. Streeting, Ozren Bogdanovic, Renata Coelho Rodrigues Noronha, Luís Adriano Santos do Nascimento, Adauto Lima Cardoso, Arthur Georges, Haoyu Cheng, Hardip R. Patel, Kishore Raj Kumar, Amali C. Mallawaarachchi, Ira W. Deveson

## Abstract

Advances in long-read sequencing (LRS) and assembly algorithms have made it possible to create highly complete genome assemblies for humans, animals and plants. However, ongoing development is needed to improve accessibility, affordability, and assembly quality and completeness. ‘Cornetto’ is a new strategy in which we use programmable selective nanopore sequencing to focus LRS data production onto the unsolved regions of a nascent assembly. This improves assembly quality and streamlines the process, both for humans and non-human vertebrates. Cornetto enables us to generate highly complete diploid human genome assemblies using only nanopore LRS data, surpassing the quality of previous efforts at a fraction of the cost. Cornetto enables genome assembly from challenging sample types like human saliva. Finally, we obtain accurate assemblies for clinically-relevant repetitive loci at the extremes of the genome, demonstrating valid approaches for genetic diagnosis in facioscapulohumeral muscular dystrophy (FSHD) and *MUC1*-autosomal dominant tubulointerstitial kidney disease (*MUC1*-ADTKD).

## INTRODUCTION

The capacity to obtain high quality and even complete telomere-to-telomere (T2T) assemblies for large eukaryotic genomes is transforming our understanding of genome architecture, variation and evolution^1–3^. The first complete T2T human genome was published in 2022, overcoming technical challenges that had left the final 8% of its sequence unsolved for two decades after the conclusion of the Human Genome Project^4^. Recent advances in the field have been driven predominantly by a handful of current US-led consortium projects, including the T2T Consortium^4^, the Human Pangenome Reference Consortium (HPRC)^5^ and Vertebrate Genome Project (VGP)^6^. These critical initiatives have led the way in molecular and computational methods development for eukaryotic genome assembly and evaluation^4-16^.

However, concentration of research and innovation within major consortium projects also reflects the high cost and high degree of technical expertise involved in producing a complete and accurate genome assembly. Current best practices call for a combination of deep Pacific Biosciences (PacBio) HiFi LRS data, coupled with Oxford Nanopore Technologies (ONT) ultra-long LRS data and Illumina HiC chromatin conformation capture, or an analogous long-range sequencing method^1^. Integration of these different data types helps to address blindspots in each. For example, the higher accuracy of PacBio HiFi is useful for untangling segmentally duplicated DNA with fine sequence differences between copies, ONT’s longer reads have capacity to span large repeats or extended regions of homozygosity, and HiC enables long-range phasing to resolve haplotypes at chromosome scale^1^. This recipe requires access to three sequencing platforms, comes with onerous sample requirements and considerable cost.

Therefore, there is a need for ongoing development to improve the affordability and accessibility of data production; increase the breadth of usable sample types and qualities; and improve data quality and assembly algorithms to better resolve the genome’s most challenging regions. Failure to address these barriers will ensure the continued exclusion of many potential research projects, cohorts and species from this new era of complete genomes.

Here we present Cornetto, a genome assembly strategy designed to meet this need. ONT’s ReadUntil or adaptive sampling functionality enables programmable selective LRS by accepting or rejecting DNA fragments, based on their sequence, in real-time^17^. This can be used to enrich genomic regions of interest, enabling targeted analysis of clinically relevant genes, for example^18–20^. We have adapted this capability to the challenge of genome assembly, integrating ReadUntil selective sequencing with the assembly process to enrich LRS data where it is most needed, thereby reducing production costs, sample requirements and improving assembly quality (**Fig1a**). Our new strategy is suitable for human and non-human genomes, diverse sample types, and accurately resolves repetitive medically-relevant loci.

**Figure 1.**
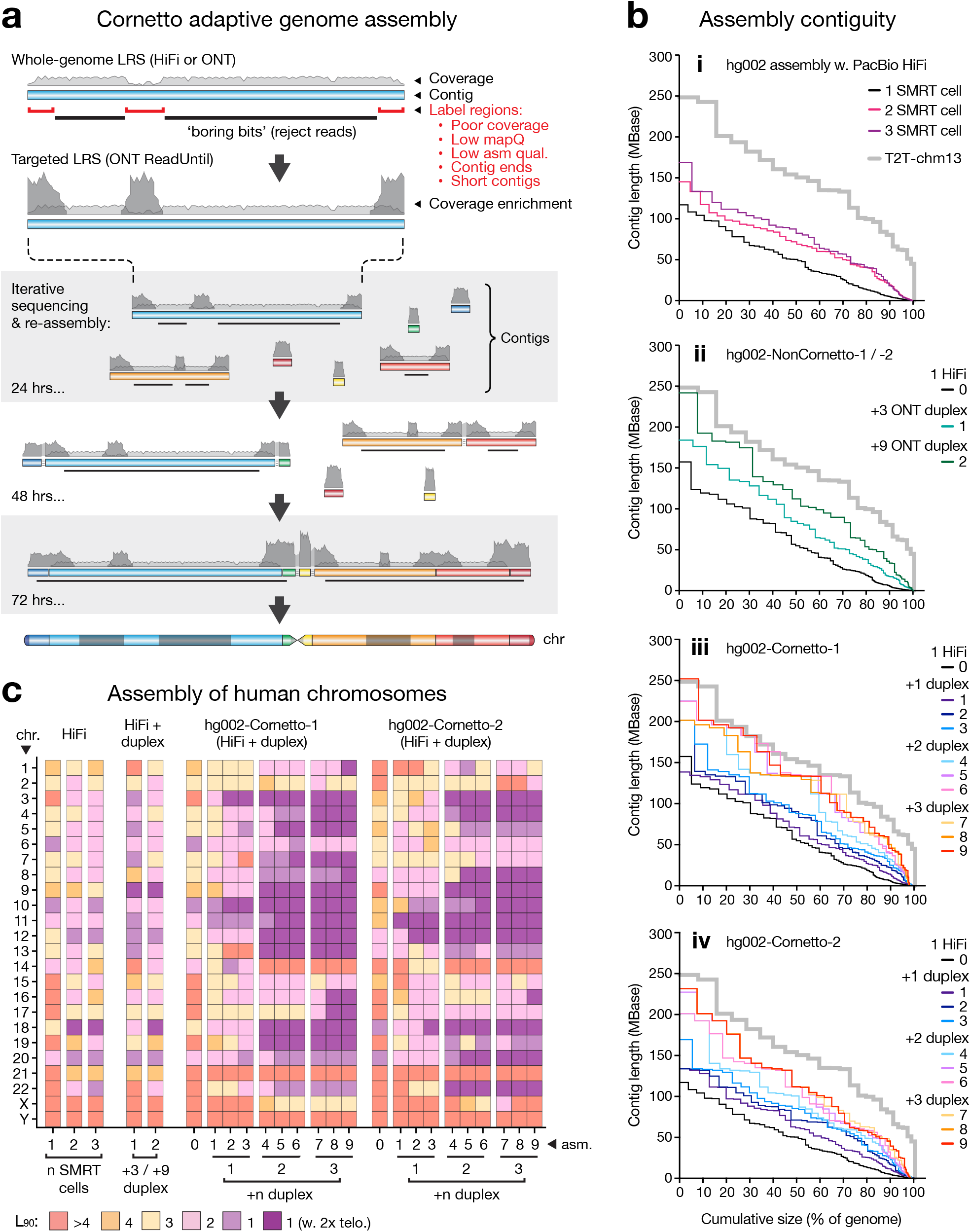
Hg002 primary assemblies. (**a**) Overview of the Cornetto method. A primary assembly generated from whole-genome LRS data (PacBio HiFi or ONT), provides the starting reference. Regions with poor coverage, mappability, assembly quality, short contigs and contig ends are identified. The remainder are considered ‘boring bits’ and labelled for rejection by ONT ReadUntil. Sequencing is paused at regular intervals (e.g. 24, 48, 72hrs) and new assembly is generated, providing an updated reference and boring bits. Sequencing is resumed after washing and re-loading flow cell. Data is focused onto an increasingly small and challenging portion of the genome. (**b**) For a given primary assembly, Nx plots show contigs sizes sorted from largest to smallest, relative to cumulative assembly size, as a percentage of human genome (3.1 Gbase). Each line shows n = 1 assembly. Assemblies were generated using LRS data from hg002. T2T-chm13 is shown for comparison (grey line). Sub-panels show: (**i**) Assemblies of HiFi data from 1, 2 or 3 SMRT cells. (**ii**) Combined data from 1 SMRT cell and 3 ONT duplex flow cells (*hg002-NonCornetto-1*); or 1 SMRT cell and 9 duplex flow cells (*hg002-NonCornetto-2*). (**iii, iv**) Two Cornetto experiments (*hg002-Cornetto-1 / -2*), each using 1 SMRT Cell and 3 duplex flow cells run sequentially with 3 cycles per cell, resulting in 9 intermediate assemblies. (**c**) For the same assemblies, tile plot shows contiguity of human chromosomes. Colour scale encodes L_90_ values: number of contigs encompassing >90% of the reference sequence for a given chromosome. Dark purple tiles show chromosomes with L_90_ = 1 and a telomere detected at each end, indicating the whole chromosome is assembled as a single primary contig.

## RESULTS

### Cornetto: iterative targeted sequencing and assembly

Most of the euchromatic human genome is relatively easy to assemble using current LRS data. For example, we sequenced DNA from the hg002 reference sample on a single PacBio SMRT cell (∼25x depth) and assembled the data with hifiasm^8^. In the resulting primary assembly, less than 10% of the genome sequence remains in contigs shorter than ∼5 Mbase (**Fig1b-i**). The assembly may be improved with further whole-genome LRS, however, there is diminishing marginal utility because most of the genome is already resolved (**Fig1b-i**).

It has been proposed that ONT ReadUntil could be used to selectively enrich regions of the genome that are difficult to assemble^21–23^ and here we have implemented this idea. Rather than defining target regions within the human reference genome based on prior knowledge, a more agnostic and efficient approach is to identify solved regions within a starting assembly of moderate quality, then program these for rejection during subsequent ONT sequencing so as to enrich for unsolved regions. This is the basis for an iterative sequencing and assembly strategy, which we nickname ‘Cornetto’ (see **Methods**).

To establish the Cornetto method, we took the hg002 primary assembly above as an initial reference, identifying short contigs (< 800 Mbase), regions adjacent to the end of a contig (within 200 kb), and regions with poor coverage, mapability or assembly quality, then labelling the remaining ∼89% of the assembly as ‘boring bits’. We then performed ONT duplex sequencing using the software ReadFish^17^ to programmably reject DNA fragments originating from any of the boring bits, in real-time (**Fig1a**). We used ONT duplex data here, because it is sufficiently similar in per-base accuracy to be co-assembled with PacBio HiFi data (**Supplementary Figure 1a-c**). The experiment was paused at regular intervals, allowing new and existing data to be aggregated and reassembled (**Fig1a**). The experiment was then resumed using the new assembly and an updated list of boring bits for rejection. The assembly and target selection processes are automated, with no manual curation required. The assembly was iteratively improved, expanding the boring bits and, thereby, focusing new data onto an increasingly small, unsolved fraction of genome (**Fig1a**).

### Human genome assembly with PacBio and ONT data

We performed two independent experiments with DNA from hg002 (*hg002-Cornetto-1* and *hg002-Cornetto-2*). Each was sequenced with one PacBio SMRT cell and three ONT duplex flow cells, which were run in succession according to the iterative Cornetto process above (3x cycles per flow cell). Comparing the primary assemblies, we observed incremental improvements in contiguity over the course of the experiments, resulting in substantial overall gains (**Fig1b-iii,iv**). For example, *hg002-Cornetto-1* was improved from 496 contigs with an N_50_ length of 54.5 Mbase and N_90_ of 6.3 Mbase to a final assembly of 130 contigs with N_50_ of 134.1 Mbase (2.5-fold improvement) and N_90_ of 50.4 Mbase (8.0-fold improvement; **Fig1b-iii,iv**; **Supplementary Data 1**). Whereas no chromosome was assembled as a single primary contig in the initial assemblies, we obtained 15 in the final assembly for *hg002-Cornetto-1* and 11 for *hg002-Cornetto-2* (**Fig1c**).

To put these results in context, we generated a matched assembly from the same PacBio HiFi data, this time augmented with three ONT duplex flow cells run without adaptive sequencing (*hg002-NonCornetto-1*; **Supplementary Figure 1a-c**). The initial assembly was improved, but the gains were small compared to those achieved using Cornetto. For example, N_50_ and N_90_ lengths were improved by 1.5-fold and 2.7-fold, respectively, in *hg002-NonCornetto-1* compared to 2.5-fold and 8-fold for *hg002-Cornetto-1* (**Fig1b-ii,c**). We further augmented the non-Cornetto assembly by adding published ONT duplex data^9^ (*hg002-NonCornetto-2*). However, even with up to nine duplex flow cells – beyond which there were no further gains – we were unable to obtain a primary assembly of comparable contiguity to *hg002-Cornetto-1* or *-2* (**Fig1b-ii,c**). Assemblies generated using Cornetto were also equivalent or superior across a range of standard quality metrics, including per-base accuracy (QV), BUSCO gene completeness, and rates of duplicated or fragmented genes (**Supplementary Data 1**). These results establish the capacity of our Cornetto strategy to efficiently harness LRS data for improved assembly of human genomes.

### Diploid human genome assemblies with ONT data

To further streamline the assembly process, we next tested the Cornetto paradigm using ONT data alone. We generated a new hg002 assembly (*hg002-Cornetto-3*), this time with data from a standard ONT flow cell (LSK114; simplex reads; **Supplementary Figure 1a-c**) augmented with a second flow cell run with Cornetto iterative sequencing (3x cycles). As above, the primary assembly was improved via Cornetto, obtaining a final N_50_ of 154.4 Mbase (1.6-fold improvement) and N_90_ of 79.1 Mbase (2.1-fold improvement; **Supplementary Figure 2a-d**; **Supplementary Data 2**).

**Figure 2.**
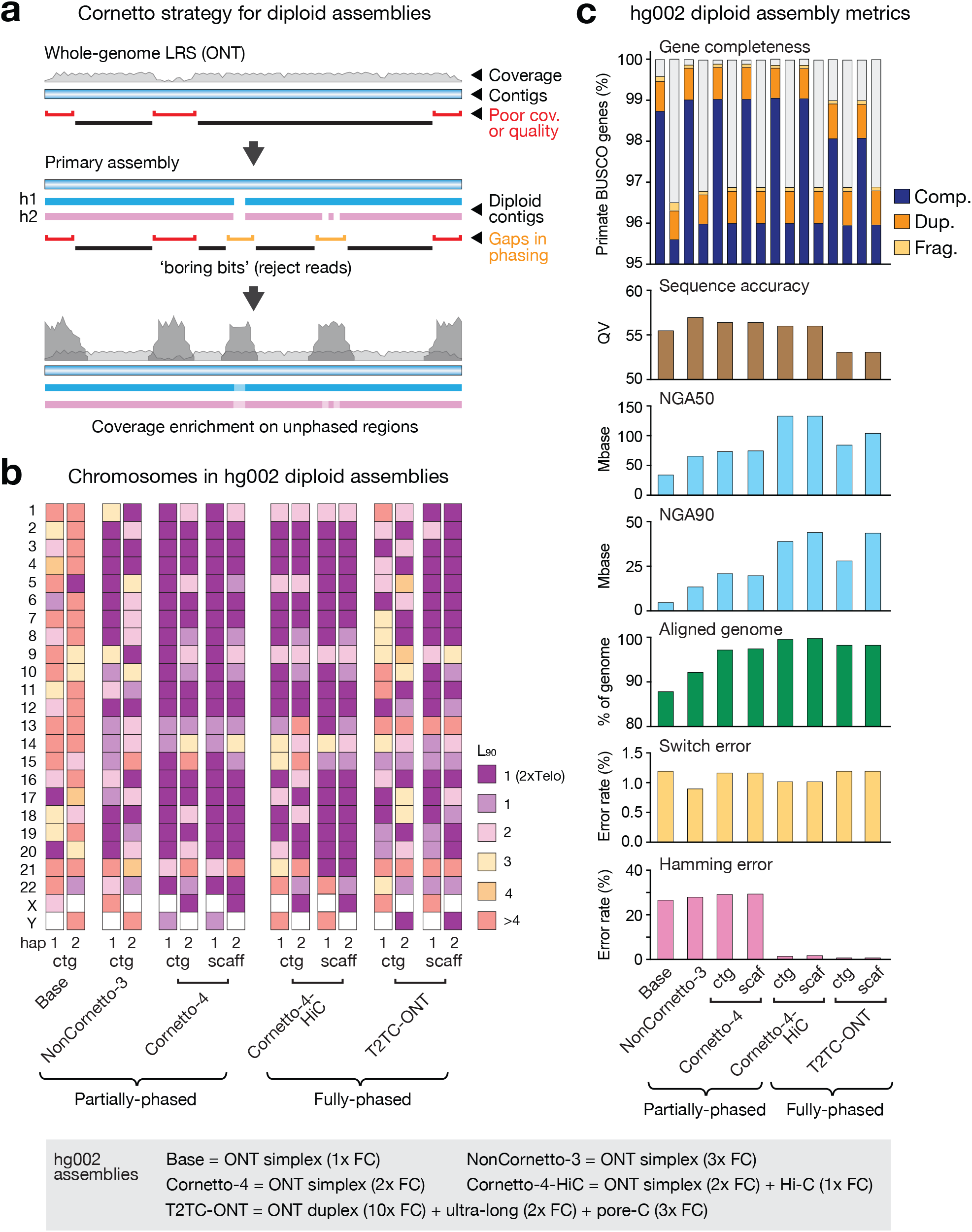
Diploid human genome assemblies for hg002. (**a**) Overview of the updated Cornetto method for improved diploid assemblies. Same process as shown in **Fig ure 1** is followed with the additional step of excluding unphased regions from the list of boring bits. These are determined by aligning all contigs from haplotype 1 & 2 to their primary assembly and identifying any region not spanned by a contig on both haplotypes. Enrichment of ONT data in these regions may help to close phase gaps. (**b**) Tile plot shows contiguity of human chromosomes in diploid assemblies for the hg002 reference sample with haplotypes plotted separately (hap1/2) and contigs vs scaffolds, where relevant. Colour scale encodes L_90_ values: number of contigs encompassing >90% of the reference sequence for a given chromosome. Dark purple tiles show chromosomes with L_90_ = 1 and a telomere detected at each contig end, indicating the whole chromosome is assembled into a single contig/scaffold. Partially-phased vs fully-phased assemblies are shown separately. Data types and sequencing resources used for each assembly are detailed in the legend below. A published ONT-only assembly from the T2T Consortium is included for comparison (T2TC-ONT). (**c**) For the same assemblies, bar charts compare assembly quality metrics: proportion of primate BUSCO genes detected as complete, duplicated, fragmented or missing; total number of gaps; sequence accuracy, as per QV values (k-mer size of 21); reference-aligned contig/scaffold N_50_ and N_90_ lengths; % of genome covered by aligned contig/scaffolds; switch errors (%); hamming errors (%). The Q100 T2T-hg002 assembly was used as a reference. All plots show data for n = 1 assembly.

Following this promising outcome, we adapted the Cornetto paradigm toward the challenge of producing diploid genome assemblies with ONT data, noting that all results reported so far refer to primary assemblies (i.e. where maternal and paternal haplotypes remain collapsed into a single, linear representation). To do so, partially-phased contigs produced by hifiasm during each Cornetto cycle were aligned to their corresponding primary assembly, to identify regions not spanned by contigs from both haplotypes (**Fig2a**). These unphased regions were excluded from the list of boring bits, which were otherwise defined as above (see **Methods**). We reasoned that the enrichment of coverage in these regions may be beneficial for closing gaps in phasing – by providing additional reads that may span a homozygous region, for example – thereby improving the resulting diploid assembly (**Fig2a**).

We applied our modified Cornetto strategy to hg002 DNA, using one standard ONT flow cell and a second run with Cornetto iterative sequencing to produce a partially-phased or ‘dual assembly’^8^ (*hg002-Cornetto-4*; i.e. where maternal and paternal haplotypes are both assembled but are expected to have occasional haplotype switches). We further supplemented this with long-range data (Hi-C), which is required to obtain a fully-phased assembly (*hg002-Cornetto-4-HiC;* i.e. in which maternal and paternal haplotypes are resolved without switches). We obtained highly complete diploid assemblies with N_90_ lengths of 59.7 Mbase for partially-phased contigs and 42.0 Mbase for fully-phased contigs (**Supplementary Figure 3a,b**; **Supplementary Data 3**). *Hg002-Cornetto-4* contained 33 partially-phased T2T chromosomes out of a possible 46, compared to just 3/46 prior to Cornetto. *Hg002-Cornetto-4-HiC* contained 32/46 chromosomes as fully-phased T2T contigs or scaffolds, including complete pairs for 14 autosomes and chrX (**Fig2b**). Alignment of *hg002-Cornetto-4* to the <u>Q100 project</u> *T2T-hg002* diploid reference, taken here as a ground truth, confirmed the assembly is free of large chromosomal misassemblies, with an aligned contig NGA_90_ length of 21.1 Mbase (**Supplementary Figure 3c,d**). The inclusion of Hi-C data improved the contig NGA_90_ length to 39.0 MBase and reduced the hamming error from 29.3% to 1.6% (**Fig2c**), as expected for a partially-vs fully-phased assembly^8^.

**Figure 3.**
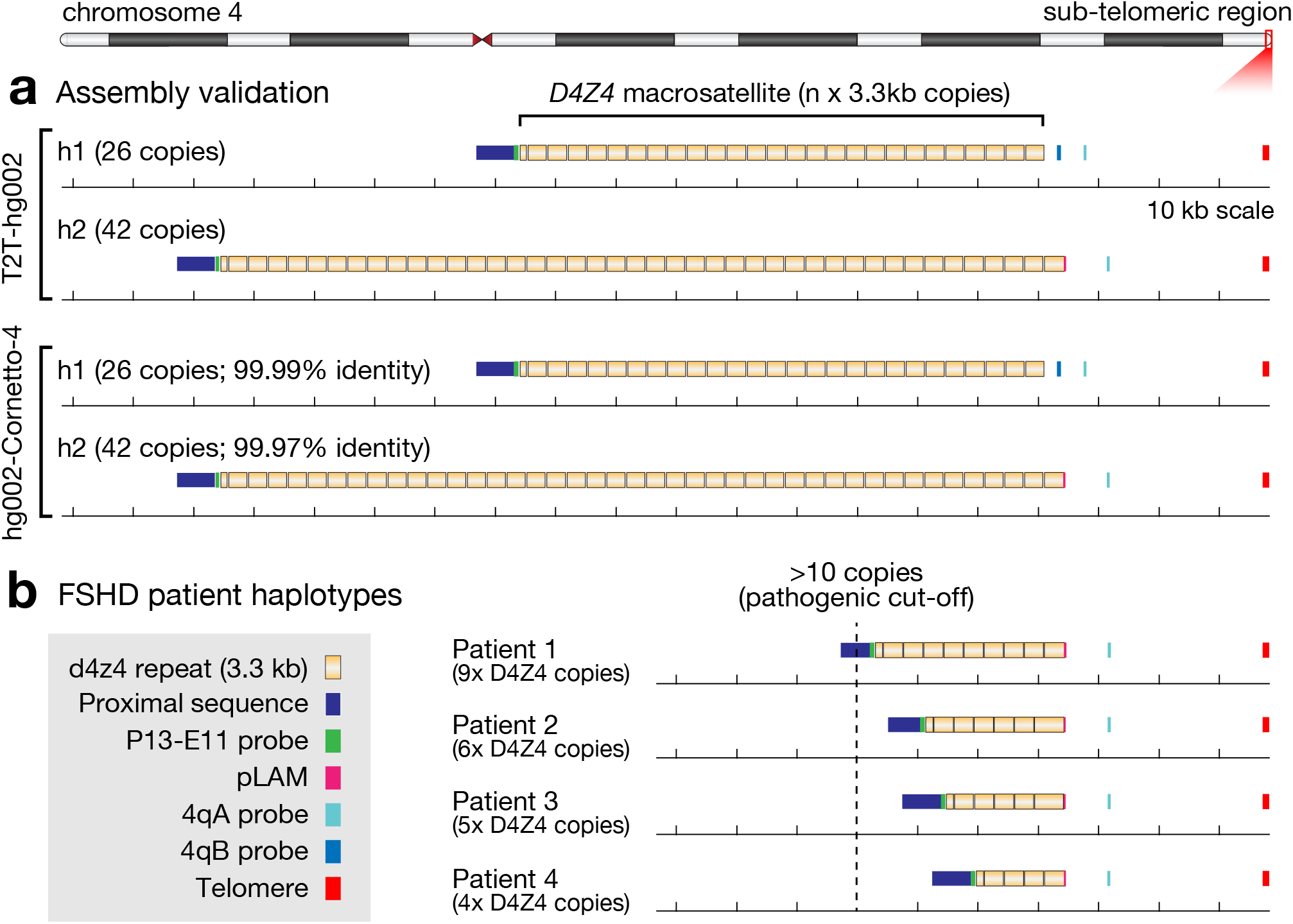
Assembly and genotyping of 4q *D4Z4* for genetic diagnosis of FSHD. (**a**) Genome browser views show annotated sequence features within the 4q subtelomeric region on each haplotype (h1 / h2) for the Q100 T2T-hg002 reference assembly (upper) and the *hg002-Cornetto-4* assembly (lower). The *D4Z4* macrosatellite repeat is annotated with recurring kb subunits in yellow. A range of other sequence features relevant for 4q *D4Z4* genotyping are shown, including markers for the permissive (4qA) and non-permissive (4qB) distal *DUX4* sequence variants (see **Supplementary Note 1**). The 4q *D4Z4* length is stated above each haplotype. Identity scores stated for hg002-Cornetto-4 were determined by aligning the entire 4q *D4Z4* sequence to the corresponding haplotype in the T2T-hg002 reference, taken as a ground truth. (**b**) Same plots as above, this time showing pathogenic haplotypes assembled using targeted ONT sequencing of the 4q *D4Z4* region for four patients with diagnostically confirmed FSHD. In each case, the 4q *D4Z4* length is shorter than 11 copies and harbours the permissive 4qA sequence variant, sufficient for a positive genetic diagnosis. Expected repeat sizes from previous genetic testing are stated for each patient (see **Supplementary Note 1**).

A recent T2T Consortium study presented a strategy for assembling diploid human genomes using data from ONT instruments alone, doing so with a combination of 50x duplex, 30x ultra-long and 50x pore-C data, utilising >15 ONT flow cells in total^9^. We next evaluated our *hg002-Cornetto-4* and *hg002-Cornetto-4-HiC* assemblies by comparison to this published, fully-phased assembly (*T2TC-ONT*). *Hg002-Cornetto-4* was slightly more complete than *T2TC-ONT* (BUSCO 99.0% vs 98.1%), showed slightly higher sequence accuracy (QV 56 vs 53), superior performance during assembly-guided structural variant detection (F1 0.952 vs 0.902), and their switch error rates were equivalent (1.2%). However, *Hg002-Cornetto-4* also showed a higher rate of hamming errors (29.3% vs 0.8%) and shorter reference-aligned contig blocks (NGA_90_ 21.1 vs 28.2 Mbase; **Fig2c**; **Supplementary Figure 3d**; **Supplementary Data 3,4**). Comparing the two fully-phased assemblies, *Hg002-Cornetto-4-HiC* was more complete, accurate and contiguous than *T2TC-ONT*, with 23 vs 13 chromosomes in complete T2T contigs and 32 vs 26 in complete T2T scaffolds (**Fig2b,c**). A small difference in hamming error (1.9 vs 0.8%) was the only standard metric we could find where *T2TC-ONT* outperformed *Hg002-Cornetto-4-HiC*. Overall, our diploid human assemblies created with Cornetto are equivalent or superior to *T2TC-ONT* on most metrics, despite using a fraction of the resources.

### Assembling medically-relevant repetitive loci

The human genome contains hundreds of analytically challenging repetitive loci with known roles in disease^13^. To illustrate the potential for Cornetto to improve inherited disease diagnosis, we next explored two such loci, which are among the most extreme known examples. In both cases, a current inability to sequence the causative locus is a barrier to effective diagnosis for its relevant disease, namely facioscapulohumeral muscular dystrophy (FSHD) and *MUC1*-autosomal dominant tubulointerstitial kidney disease (*MUC1*-ADTKD). The unmet needs and challenges involved are described in more detail in **Supplementary Notes 1** and **2**, respectively.

FSHD is a progressive myopathy resulting from aberrant expression of the *DUX4* gene residing within the 4q *D4Z4* macrosatellite repeat, a polymorphic *n* x 3.3kb tandem repeat in the sub-telomeric region of chr4q^24^. The repeat typically ranges in size from ∼11–100 copies^24^. FSHD most commonly presents in individuals with a contracted 4q *D4Z4* haplotype (<10 copies), which must also harbour a permissive sequence variant (4qA) following the distal-most *DUX4* copy^24^. To assess our capacity to accurately assemble this locus, we extracted the sub-telomeric region from both copies of chr4q in the *hg002-Cornetto-4* assembly, annotated known sequence features relevant to FSHD, then compared them to the equivalent regions of the *T2T-hg002* reference, again taken as ground truth. In both assemblies we identified one *D4Z4* haplotype at 42 copies in length (∼139 kb) of subtype 4qA and a second at 26 copies (∼86 kb) of subtype 4qB (**Fig3a**). Aligning corresponding haplotypes between the two assemblies, we observed 99.99% and 99.97% sequence concordance across entire 4q *D4Z4* regions, with 1-2bp insertions/deletions accounting for most of the discrepancies (31/35; **Fig3a**). We next performed targeted ONT sequencing and assembly of this region in four patients with diagnostically confirmed FSHD. In each case we identified one *D4Z4* haplotype of the permissive 4qA sub-type with fewer than 10 copies, sufficient for a positive diagnosis, and observed repeat lengths that were concordant with previous molecular genetic testing (**Fig3b**; see **Supplementary Note 1**).

ADTKD is a chronic kidney disease typically caused by variants in one of four genes, *UMOD, MUC1, REN* and *HNF1B*^25^. *MUC1* is thought to account for around ∼20% of cases, however, diagnosis of *MUC1*-ADTKD is obscured by technical challenges in resolving this gene. *MUC1* contains a *n* x 60bp variable number tandem repeat region (VNTR)^26^. This is highly polymorphic, varying in length (20–125 copies per haplotype) and differing in the composition of imperfect sequence subunits within and between individuals^5,26^. Duplication of a cytosine (dupC) within this VNTR, which results in a frameshifting variant, has been identified as the predominant cause of *MUC1*-ADTKD^27^ (**Fig4a**). We identified both copies of the *MUC1* locus within our *hg002-Cornetto-4* assembly, annotated the VNTR region for known and novel 60bp subunits, and compared them to *T2T-hg002* (**Fig4b**). In both assemblies we identified one VNTR haplotype with 65 x 60bp copies (∼3.9 kb) and one with 78 x 60bp copies (∼4.7 kb). The composition and order of VNTR subunits was matched and, aligning corresponding haplotypes between assemblies, we observed perfect sequence concordance across the entire VNTR region (**Fig4b**). We next performed targeted ONT sequencing and *MUC1* assembly in ten patients with diagnostically confirmed *MUC1*-ADTKD. Across 11 individuals (including *hg002-Cornetto-4*), we observed 20 unique VNTR haplotypes which ranged in size from 40–83 copies, with no individuals sharing the same pair of haplotypes (**Fig4b**). In each patient (but not hg002) we identified a single dupC frameshift variant within the VNTR occurring on a single haplotype, sufficient for a positive diagnosis. Notably, 8/10 pathogenic haplotypes were unique, implying frequent independent origins of the dupC variant (**Fig4b**; see **Supplementary Note 2**). Overall, these results establish the capacity to accurately assemble both the 4q *D4Z4* and *MUC1* loci, providing viable new avenues to improve the genetic diagnosis of FSHD and ADTKD.

**Figure 4.**
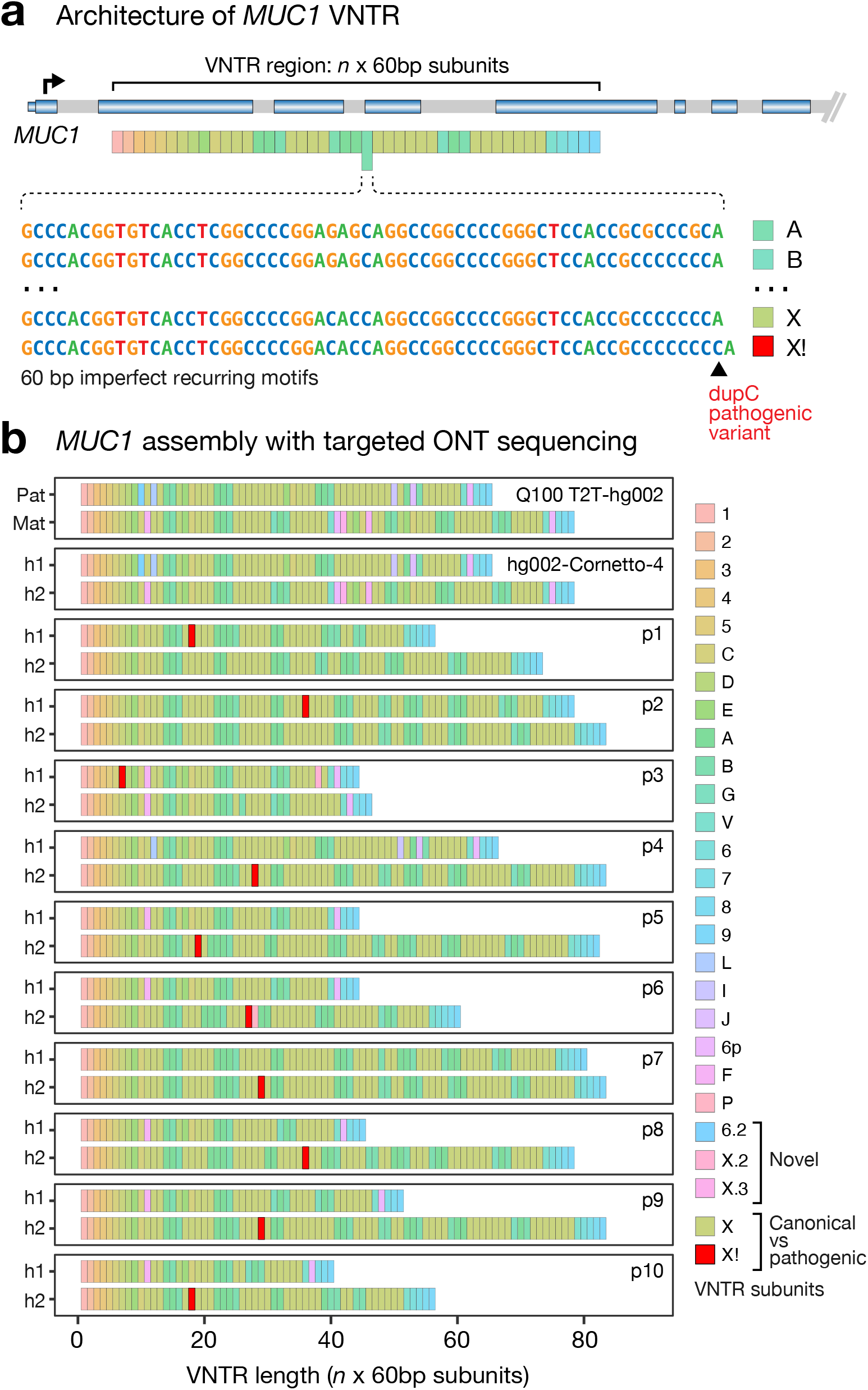
Assembly and genotyping of *MUC1* for genetic diagnosis of ADTKD. (**a**) Schematic of *MUC1* variable number tandem repeat region (VNTR) and known genetic basis for *MUC1*-ADTKD. Briefly, *MUC1* contains a large VNTR comprising recurring imperfect 60bp subunits, which varies in length and sequence composition within and between individuals. Duplication of a cytosine (dupC) within the VNTR, resulting in a frame-shift causes *MUC1*-ADTKD (see **Supplementary Note 2**). (**b**) Tile-bar plots show the VNTR length and subunit composition identified for each *MUC1* haplotype (h1 / h2) for the Q100 T2T-hg002 reference (upper) and hg002-Cornetto-4 assembly, which show identical length and sequence composition between corresponding haplotypes. Below are VNTR haplotypes assembled via targeted ONT sequencing in ten patients with diagnostically confirmed *MUC1*-ADTKD. Different coloured tiles indicate known and novel 60bp sequence subunits, with the known dupC pathogenic subunit (X!) shown in red. A single X! subunit was identified on one haplotype in each patient, sufficient for a positive diagnosis (see **Supplementary Note 2**).

### Genome assemblies from human saliva

Another benefit of Cornetto is to enable production of high quality assemblies from challenging and/or limited sample types. Human saliva is one such sample type where there would be significant utility. Saliva is more easily accessible than blood or other human tissues and can be collected, shipped and stored at room temperature. This can be advantageous in some clinical contexts, for field studies in remote communities^28^ or even for direct-to-consumer genomics^29^. However, saliva is less amenable than blood to extraction of high-molecular weight (HMW) DNA; is not compatible with ONT’s ultra-long protocol, nor HiC or other related long-range methods; and often suffers from relatively high levels of non-human DNA contamination^30^. Given these challenges, we are not aware of previous attempts to assemble a human genome from saliva.

We collected saliva from a male and female participant, extracted HMW DNA, then conducted Cornetto sequencing and assembly on each. We tested a combined PacBio HiFi and ONT duplex approach (*saliva-A-Cornetto-1*; *saliva-B-Cornetto-1*) and an ONT-only approach (*saliva-A-Cornetto-2*; *saliva-B-Cornetto-2*). Because non-human DNA was present (**Fig5a**), we additionally included non-human contigs identified in the initial assemblies in the target list for rejection, selecting against further contamination (see **Methods**). Both Cornetto approaches yielded high-quality genome assemblies with improved contiguity and completeness relative to their starting assemblies (**Fig5b-d**; **Supplementary Figure 4a-c**; **Supplementary Data 5**). The improvements were particularly pronounced for ONT-only diploid assemblies *saliva-A-Cornetto-2* and *saliva-B-Cornetto-2*, for which we obtained final contig N_90_ lengths of 46.5 Mbase (15-fold) and 50.1 Mbase (27-fold), and 27/46 and 26/46 complete chromosomes, respectively (**Fig5b,c**). Despite the use of saliva as input material, these results are comparable to the *hg002-Cornetto-4* and *T2TC-ONT-only* assemblies above.

**Figure 5.**
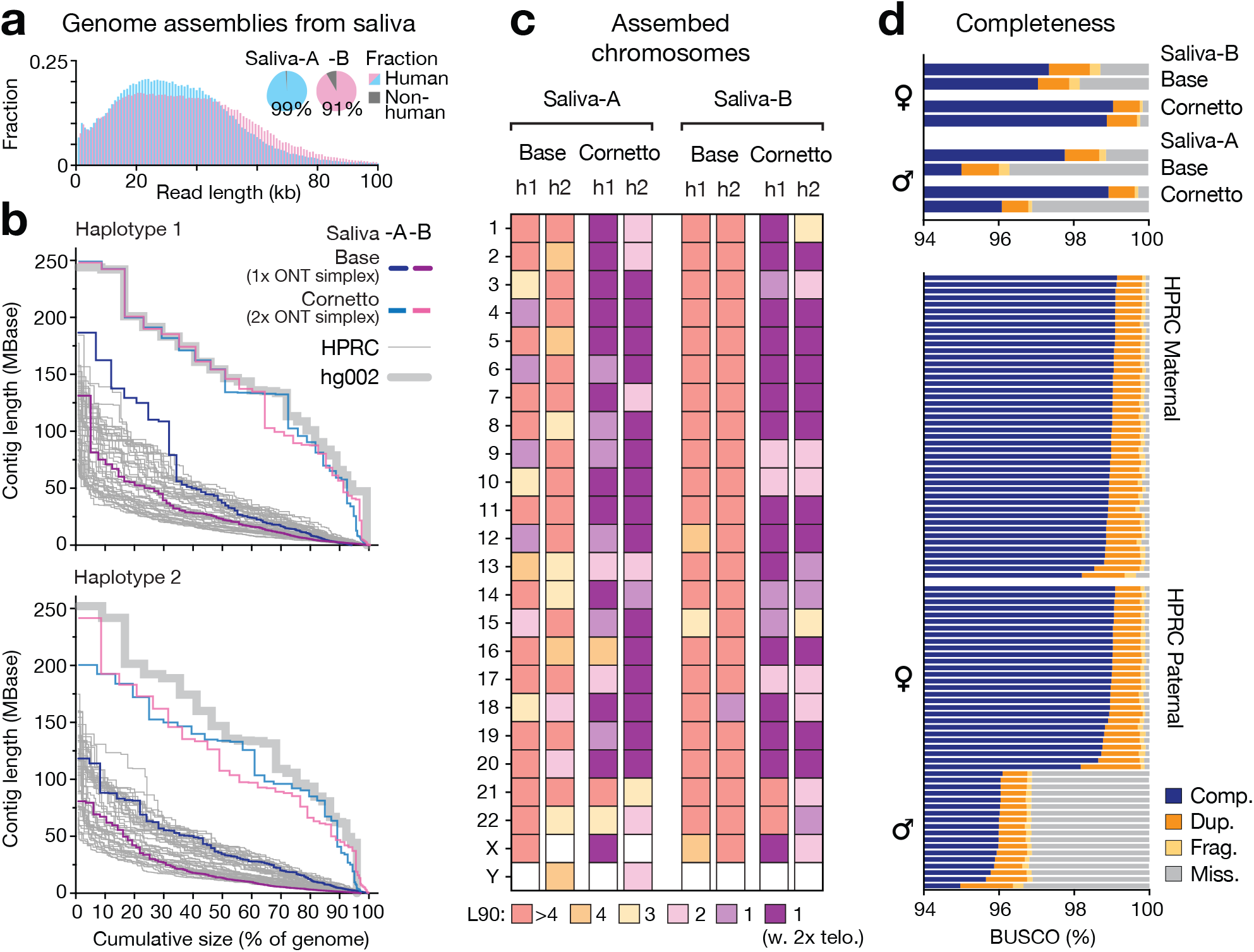
Genome assemblies from human saliva. (**a**) Histograms show read length profiles and pie charts show proportion of non-human reads from standard ONT sequencing on saliva samples from two participants: Saliva-A (male) and Saliva-B (female). (**b**) Nx plot shows contig sizes sorted from largest to smallest, relative to cumulative assembly size, as a percentage of the human genome size (3.1 Gbase). For each participant, assembly generated using ONT data from one flow cell (base) then augmented with a second flow cell run with Cornetto are shown (Saliva-A-Cornetto-2, Saliva-B-Cornetto-2). Non-human reads were excluded prior to assembly. For comparison, thin grey lines show contig sizes for all assemblies released in the first phase of the HPRC (*n* = 47) and thick grey lines show the Q100 T2T-hg002 assembly. Diploid haplotypes for each assembly are divided between the two plots. (**c**) For the same assemblies, tile plot shows contiguity of human chromosomes. Colour scale encodes L_90_ values: number of contigs encompassing >90% of the reference sequence for a given chromosome. Dark purple tiles show chromosomes with L_90_ = 1 and a telomere detected at each contig end, indicating the whole chromosome is assembled as a single contig. (**d**) For the same assemblies as **b**, stacked bar charts show the proportion of BUSCO genes detected as complete, duplicated, fragmented or missing. Haplotype groups containing the Y-chromosome sequence have a larger proportion of missing genes (designated with male marker symbol). All plots show data for n = 1 assembly.

For further context, we compared our saliva assemblies to 47 assemblies released in the first phase of the HPRC, which were assembled with hifiasm using HiFi and trio sequencing data^5^. *Saliva-A-Cornetto-2* and *Saliva-B-Cornetto-2* exhibited comparable or superior BUSCO gene completeness and substantially better contiguity than any HPRC assembly, although we note the saliva assemblies are only partially-phased (**Fig5b,d**). Although assembly accuracy cannot be directly measured, as no ground truth is available, the results presented above for *hg002-Cornetto-4* imply comparability with HPRC assemblies on these parameters. In summary, Cornetto can be used to obtain highly complete assemblies from human saliva, which are in line with (or surpass) quality standards at the leading edge of the genomics field.

### Genome assemblies for non-human vertebrates

Cornetto is sequence-agnostic and does not rely on any prior knowledge of the genome being assembled. In theory, the method is suitable for any species. To establish this, we next assembled genomes for a selection of non-human vertebrates from diverse lineages, prioritised for their salience in research and conservation. The critically endangered orange-bellied parrot (*Neophema chrysogaster*) and endangered western saw-shelled turtle (*Myuchelys bellii*) were assembled using only ONT data, while Gould’s petrel (*Pterodroma leucoptera*; a threatened seabird) and the redstriped eartheater cichlid (*Geophagus surinamensis*; an Amazonian fish) were assembled with PacBio HiFi and ONT duplex data (see **Supplementary Note 3**).

For each species, Cornetto delivered improvements in genome assembly outcomes, compared to base assemblies generated with standard LRS data (**Fig6a-c**; **Supplementary Data 6**). For example, we obtained a 3.9-fold increase in contig length N_50_ for the petrel primary assembly and an increase in the number of chromosomes assembled as single primary contigs from 6 to 27 (**Fig6a-c**). ONT-only diploid assemblies were also strongly improved. In the turtle genome, for example, each haplotype harboured ten complete chromosomes, including examples of both macro and microchromosomes, and complete single copies for 99.8% and 99.6% of BUSCO genes (**Fig6b,c**). Importantly, these improvements were obtained despite wide variation in genome sizes, architecture, depth of starting LRS data and initial assembly quality (see **Supplementary Note 3**). For example, the cichlid genome was initially sequenced to 78x depth with HiFi data, yielding a base assembly of 142 contigs. Cornetto still found room for improvement, reducing the number of primary contigs to 75, with a 3.8-fold improvement in contig length N_90_ and 12 chromosomes added (**Fig6a**; **Supplementary Figure 5a**). In contrast, the parrot genome was sequenced to 35x depth with standard ONT data on DNA extracted from a frozen liver sample, yielding a base assembly of >2000 contigs. With a single additional ONT flow cell run with Cornetto we were able to obtain a diploid assembly with a 4.5-fold increase in contig length N_50_ and a 4.6% increase in BUSCO completeness (92.1% vs 96.7%; **Fig6a,c**; **Supplementary Figure 5b**). Notably, our updated assembly for the orange bellied parrot contained an 80kb region with three identifiable major histocompatibility complex genes (*MHCI, II-A, II-B*), which are of critical importance for understanding the decline of immunogenetic diversity that threatens the survival of the species, whereas a recent assembly created with HiFi and HiC data was unable to resolve the *MHC* region^31^ (see **Supplementary Note 3**). In summary, this establishes the suitability of our Cornetto assembly paradigm for assembling diverse non-human genomes.

**Figure 6.**
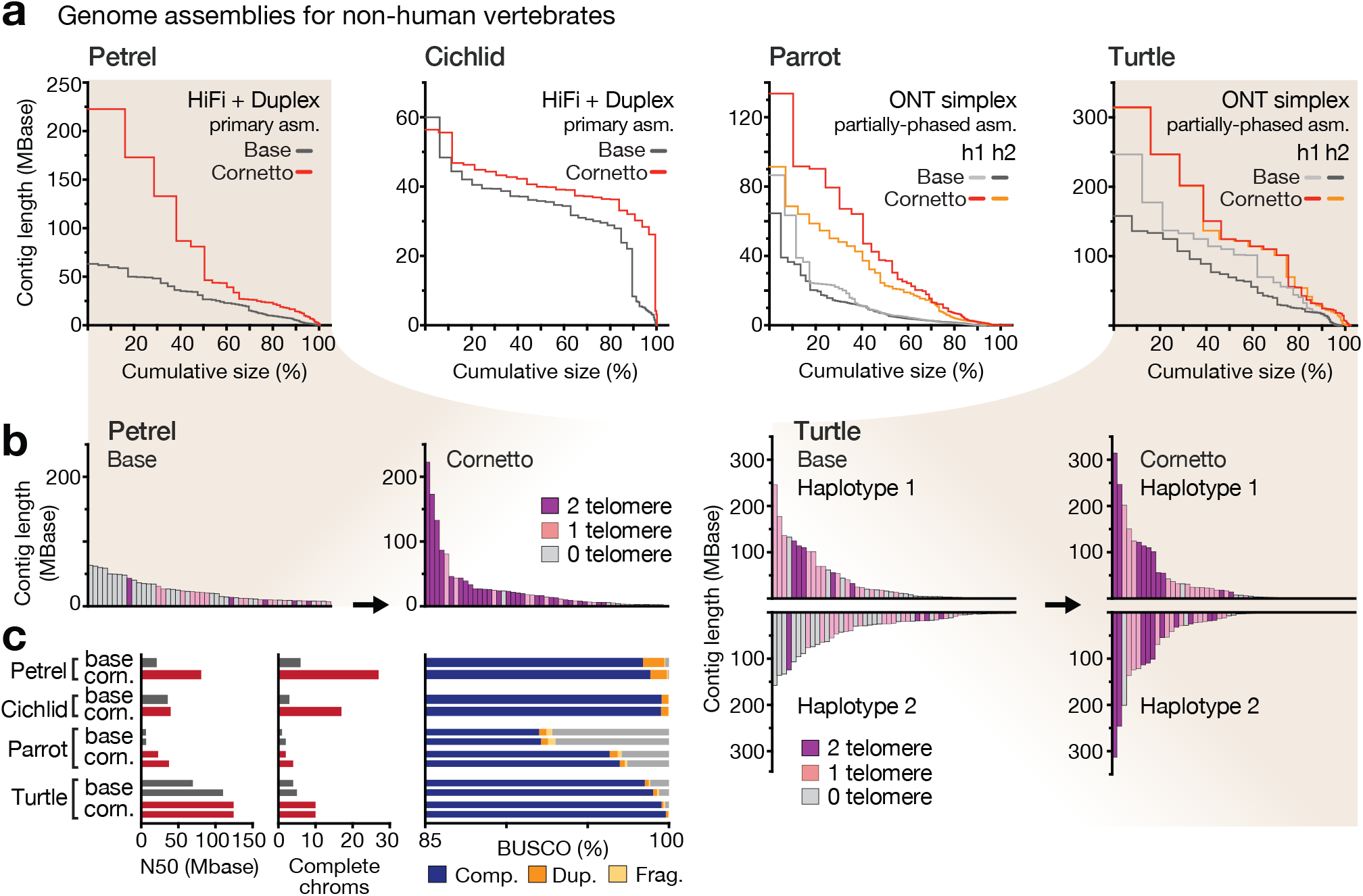
Genome assemblies for non-human vertebrates. (**a**) Nx plot shows contig sizes sorted from largest to smallest, relative to cumulative assembly size, as a percentage of the haploid genome size for each species. From left to right the plots show genome assemblies for: Gould’s petrel (*Pterodroma leucoptera*); redstriped eartheater cichlid (*Geophagus surinamensis*); orange-bellied parrot (*Neophema chrysogaster*); western saw-shelled turtle (*Myuchelys bellii*; see **Supplementary Note 3**). The petrel and cichlid were assembled with PacBio HiFi (1 SMRT cell; base) plus ONT duplex data (2 duplex cells; cornetto). The parrot and turtle were assembled using ONT data, one or two standard flow cells (base) plus another with adaptive sequencing (Cornetto; simplex reads; LSK114). For ONT-only experiments, diploid assemblies were generated and haplotypes are plotted separately. (**b**) For the petrel and turtle assemblies above, bar plots show sizes of the fifty largest contigs in descending order, coloured according to presence of telomere sequences at contigs ends (both ends = purple; one end = pink). Equivalent plots for the cichlid and parrot are shown in **Supplementary Figure 5.** (**c**) Bar plots show standard quality metrics for the same assemblies as in **a**. From left to right, these are: contig N_50_ lengths in Mbases; number of complete chromosomes (>1 Mbase and telomere detected at both ends); proportion of BUSCO genes detected as complete, duplicated, fragmented or missing. All plots show data for n = 1 assembly.

## DISCUSSION

Cornetto is a new approach to genome assembly, which is applicable to both human and non-human genomes. We have used Cornetto to generate diploid human genome assemblies for hg002 (using cultured cells) and from saliva samples. These are notable for their completeness, accuracy, and for the modest resources used in their creation, including compute resources (**Supplementary Data 7**). We obtained a high-quality human genome assembly (*hg002-Cornetto-4*) from a single ONT library sequenced on two flow cells on a portable ‘P2 Solo’ device (roughly the size of a brick), without the need for PacBio or Illumina data, which require large instruments with substantial capital costs. Similarly, *hg002-Cornetto-4* did not use ONT ultra-long or pore-C methods, which are sensitive preparations requiring access to cultured cells or freshly drawn blood. We have shown that Cornetto is compatible with ONT’s highly accurate duplex data type. However, this was not needed to produce our best assemblies, nor were computationally expensive error correction methods using deep learning (HERRO^32^ and Dorado Correct). Overall, Cornetto improves genome assembly outcomes, while streamlining the process and enhancing accessibility.

Although high-quality assemblies can be obtained using only LRS data, these are partially-phased or ‘dual assemblies’. Full resolution of parental haplotypes, at chromosome scale, requires additional data from a long-range method, such as Hi-C, which we show here can be co-assembled with Cornetto-enriched LRS data. Using this approach (*hg002-Cornetto-4-HiC*), we obtained fully-phased T2T contigs or scaffolds for 32/46 chromosomes. Notably, this included the hemizygous chrX and the acrocentric chromosomes chr13 (one haplotype) and chr21 (both haplotypes). Acrocentric chromosomes (chr13, chr14, chr15, chr21, chr22) are characterised by repetitive ribosomal DNA arrays, which pose significant challenges, and none were fully assembled in the T2TC-ONT assembly used for comparison^1,4^. Therefore, the combination of Cornetto-enriched LRS and long-range data appears promising for tackling acrocentric chromosomes; although there is much room for improvement. Recent improvements to the hifiasm^33^ software have been critical for Cornetto and we anticipate future updates may help to resolve the remaining genome regions. We also note that both ONT ultra-long and pore-C preparations are compatible with ONT selective sequencing, so may be successfully integrated with Cornetto in the future. Another intended improvement is to enable iterative updating of the genome assembly and its associated ‘boring bits’ in real-time during ONT sequencing. Currently, re-assembly is performed at experimental pause-points, requiring around ∼4-6 hours. Significant software acceleration is needed to enable real-time assembly.

Cornetto works by selectively enriching LRS data onto unsolved regions of a nascent assembly. This concept has been proposed previously^21,23^ and tested with some success in *Arabidopsis*^22^. Cornetto is the first streamlined, generic implementation and rigorous evaluation of this approach. An alternative approach would be to select static, predefined target regions within the human reference genome, which are known to be challenging.

However, this requires a high quality existing reference genome and prior knowledge to define target regions and is therefore unsuitable for most non-human species. The optimal target space may also differ between individuals based on their specific genome architectures, being strongly influenced by features such as repeat lengths and homozygous regions, which vary between individuals. The optimal target space may also differ depending on the nature of available data (read length, depth, accuracy). We therefore believe the most efficient approach is simply to select for any region that has not yet been confidently assembled, defining this in the context of a specific genome and all available data.

However, there are scenarios where targeting static reference regions may be preferable. For example, we used this approach to specifically enrich 4q *D4Z4* and *MUC1* to efficiently and reproducibly assemble these loci in patient samples. An inability to effectively sequence these loci is a major barrier to genetic diagnosis of FSHD and *MUC1-*ADTKD, respectively (see **Supplementary Notes 1** & **2**). There is also much that remains to be learned about these diseases. For example, we show here that the dupC variant known to cause *MUC1*-ADTKD has reproducibly, but independently, emerged within different families or ancestry groups, reflected in the lack of shared pathogenic haplotypes between patients. Although dupC is overwhelmingly the most common causative variant reported in *MUC1-*ADTKD^34^, this is heavily biased by the available diagnostic technique being targeted to this specific variant^26^. We anticipate that the accurate, haplotype-resolved assembly of the *MUC1* VNTR in currently undiagnosed patients will reveal additional pathogenic variants.

There is clear clinical utility in being able to genotype these and other challenging medically-relevant loci from patient saliva, but developing the capacity to assemble highly complete human genomes from saliva is an even greater priority for our Australian Indigenous genomics research program^28^. Collection of blood during visits to remote Aboriginal communities is impractical and blood is culturally sensitive, thus limiting us to work with saliva^28^. We hope that our demonstration of scalable, affordable genome assembly from saliva – with assembly quality at least on par with the HPRC – may open the door to even greater diversity and inclusion in the growing pangenome field^35^. Applications for Cornetto are not limited to human genomics, as we have shown here by producing highly complete genomes for diverse non-human vertebrates, including three critically endangered, endangered or threatened endemic Australian species. By improving costs, sample input requirements and providing higher quality reference genomes than previously possible, we hope Cornetto will empower other genomically-informed conservation initiatives, similar to the orange-bellied parrot^31^, described above. We provide all laboratory and computational methods for Cornetto assembly as a free, open source resource to streamline, improve and democratise genome assembly.

## METHODS

### Ethics and inclusion

The research complies with all relevant ethical regulations. The study design and conduct complied with all relevant regulations regarding the use of human study participants and was conducted in accordance to the criteria set by the Declaration of Helsinki. Saliva samples were collected from individuals not known to be affected by inherited disease under Human Research Ethics Committee (HREC) approval 95179 (St Vincent’s Hospital Human Research Ethics Committee). Whole blood was collected from patients with FSHD under HREC/2019/ETH12538 (St Vincent’s Hospital Human Research Ethics Committee) and patients with *MUC1*- ADTKD under HREC/18/RPAH/726 (Royal Prince Alfed Hospital Human Research Ethics Committee), HREC/83945/RCHM-2022 (Royal Children’s Hospital Melbourne Human Research Ethics Committee) and HREC/16/MH/251 (Monash Health Human Research Ethics Committee). Wherever sex, age or other potentially identifiable data is reported, this was done so with appropriate consent. Relevant approvals for non-human vertebrates are provided in **Supplementary Note 3**.

### Cornetto iterative targeted sequencing and assembly method

Cornetto is a new experimental paradigm in which the genome assembly process is integrated with ONT ReadUntil programmable selective sequencing (also known as ‘adaptive sampling’). Cornetto encompasses both laboratory and computational protocols, all of which are described here. Computational steps are executed using the open source Cornetto software package (https://github.com/hasindu2008/cornetto) in addition to other third-party open source software. Most importantly, we have used the excellent software hifiasm^8^ for generating assemblies and Readfish^17^ for executing ONT targeted sequencing. Hifiasm can be run on the user’s preferred machine using commands provided in **Supplementary Note 4** or Cornetto online documentation. ReadFish must be executed on the computer that runs ONT sequencing experiments, using reference files generated by Cornetto. Cornetto is also compatible, in principle, with ONT’s in-built ‘adaptive sampling’ application, which is configured within MinKNOW, and with alternative assembly software, however, these have not been extensively tested.

The Cornetto method starts with a primary assembly of moderate quality, generated with PacBio HiFi or ONT LRS data. For a human genome, LRS data from a single PacBio SMRT cell or a single ONT PromethION flow cell typically yields a suitable base assembly. Our recommended protocols for HMW DNA extraction, DNA shearing, size selection and library preparation, for both PacBio and ONT, are outlined in detail below. A human base assembly is created from the initial LRS data using hifiasm (versions and commands in **Supplementary Note 4**).

After generating a base assembly, the available LRS data is realigned to this assembly (using minimap2^36^) to assess regional coverage and mappability. Cornetto software is then used to identify assembly regions that are not yet confidently resolved. These are defined as: extended regions of low or high coverage depth (< 40% or > 250% of genome average); low mappability (where coverage depth in uniquely aligned MapQ20+ reads is < 40% of total mean coverage); low assembly quality (‘lowQ’ regions of > 8 Kbase output by hifiasm); short primary contigs (< 800Mbase) and; regions adjacent to the end of a primary contig (within 200 kb). For diploid assemblies, Cornetto additionally identifies unphased regions in the assembly. To do so, all contigs from haplotype 1 and haplotype 2 (output by default by hifiasm) are aligned to the primary assembly to identify any region that is not spanned by a contig in both haplotypes. These unphased regions are combined with the other labelled regions above; coordinates of all regions are then extended with 40 kb buffers in either direction before merging all overlapping and adjacent features (within 200 kb). All sequences outside of these merged regions, which typically encompass ∼10-20% of the genome, are considered to be already confidently assembled and will not benefit from additional LRS data. Hence these regions are considered ‘boring bits’ and printed to a standard coordinate file ‘boringbits.bed’ and corresponding file ‘boringbits.txt’, which is used for ReadFish configuration. Commands to execute this process are provided in **Supplementary Note 4**.

Next, the user should perform ONT sequencing with ReadFish (or the ONT adaptive sampling app) configured to reject reads originating from any of the boring bits within the initial assembly. The relevant base assembly is provided as a reference, after indexing with minimap2. Commands, software versions and a template with parameters for ReadFish configuration are provided in **Supplementary Note 4**.

After configuring and launching ReadFish (or the ONT adaptive sampling app within MinKNOW) the user should load their sequencing library onto their ONT flow cell and initiate the sequencing experiment. When starting with a base assembly of PacBio HiFi data, we recommend to run ONT ‘duplex’ sequencing because HiFi and duplex data are sufficiently similar in accuracy to be co-mingled during assembly. If starting from a base assembly generated with ONT simplex data, the user should continue with ONT simplex data. Protocols used during our study for both ONT sequencing options, including basecalling, are outlined in detail below.

To maximise yields during ONT sequencing, it is standard practice to pause the sequencing process at regular intervals, wash the flow cell with a nuclease solution, reload with fresh library, then resume sequencing. During a Cornetto experiment, the user should take advantage of these pause points in order to update their assembly and regenerate the boringbits reference files. When updating the reference files at each pause point, Cornetto uses the same rules as above, with the exception that poor coverage and mappability rules are not applied beyond the first cycle (because adaptive sequencing introduces uneven coverage which would conflict with these rules). When working with saliva samples, nonhuman contigs may also be added to the list of boringbits for rejection at this point (see below). This process serves to focus ongoing data generation onto an increasingly small and challenging portion of the genome that remains unassembled. We typically perform 3 or 4 Cornetto cycles for a single ONT flow cell, with pause points at ∼24 hr, ∼48 hr and ∼72 hr (if the flow cell remains viable). Each new assembly should be generated by aggregating all existing and new data (see **Supplementary Note 4**). At the end of the process, the user should obtain a final assembly encompassing LRS data from all previous steps. The Cornetto software also provides a simple wrapper script used to evaluate this assembly with a range of standard metrics. The evaluation process run by Cornetto and employed during this study are outlined in detail below with software versions and commands in **Supplementary Note 4**. Specifics of the process used for FSHD and ADTKD patients are outlined in **Supplementary Note 1** and **2**, respectively. Specifics of the process used for non-human vertebrates are outlined in **Supplementary Note 3**.

### High-molecular weight DNA extractions

High–molecular weight (HMW) genomic DNA was extracted from cultured cells from human reference sample hg002, obtained from the Coriell Institute for Medical Research (B-Lymphocyte cell line GM24385), and peripheral blood samples from patients with FSHD and ADTKD, using the PacBio Nanobind CBB kit (102-301-900) or PacBio PanDNA kit (103-260-000) according to the manufacturer’s protocol. For saliva experiments, ∼5mL of saliva was collected from healthy donors using Oragene self-collection kits (DNA Genotek) and stored at room temperature. Saliva was extracted using the PacBio Nanobind CBB kit with an optimised protocol that is now supported by the manufacturer (103-544-000). For experiments involving non-human vertebrates, HMW DNA was extracted from blood for the petrel, cichlid and turtle and from snap frozen liver tissue for the orange bellied parrot (see **Supplementary Note 3**). For blood samples, approximately 20-70 ul was used as input (or equivalent for ethanol stored blood samples) and these were extracted as per the PacBio Nanobind protocol for nucleated blood. For the liver tissue, 20 mg was used as input and this was extracted using the PacBio Nanobind kit standard Dounce homogenizer protocol. QC checks were performed on all extracted DNA samples using a ThermoFisher NanoDrop (purity), ThermoFisher Qubit (DNA concentration) and Agilent Femto Pulse (Genomic DNA 165 kb Kit; DNA fragment size profiles).

### Long-read sequencing methods

For PacBio sequencing experiments, DNA samples were sheared to average fragment lengths of 15–24 Kb using a Diagenode Megaruptor 3 with Shearing Kit at a speed of 30 or 31. Sheared DNA was cleaned, concentrated, then subject to PacBio SMRTbell library preparation, all according to the manufacturer’s protocol. Prepped libraries were size selected with a 35% v/v dilution of AMPure PB beads at a ratio of 2.9x (i.e. 50 ul of sample : 145 ul 35% beads) or using a PippinHT (Sage Science) with a 10 kb cut-off, followed by ABC loading procedure and sequencing on a PacBio Revio instrument with 30 hour movie time.

For ONT sequencing experiments, DNA samples were sheared to average fragment lengths of 43–66 Kb using a Diagenode Megaruptor 3 with Shearing Kit and a shearing speed of 27. Sheared samples were treated with PacBio Short-Read Eliminator kit to deplete fragments < 10 kb. ONT libraries were then prepared from ∼5–9 µg of sheared HMW genomic DNA using a ligation prep (SQK-LSK114). The resulting libraries were loaded on an ONT PromethION R10.4.1 flow cell (FLO-PRO114M) or a PromethION high-duplex flow cell (FLO-PRO114HD) and sequenced on a PromethION instrument (P2 Solo or P48). Sequencing experiments were typically run for 72 hours, with washes (EXP-WSH004) and library reloading performed at approximately 24 and 48-hour time points. Where flow cells were still viable, an additional wash was performed at 72 hours, followed by a further 24 hr runtime. For Cornetto experiments, live target selection/rejection was executed during the run by the ReadFish^17^ software package with commands and configuration parameters in **Supplementary Note 4**.

Raw ONT sequencing data was converted from POD5 to BLOW5 format^37^ in real-time during sequencing. At pause-points during sequencing, or after completion of a run, data was base-called using slow5-dorado (v0.8.3; https://github.com/hiruna72/slow5-dorado) with a recent ‘super-accuracy’ model (dna_r10.4.1_e8.2_400bps_sup@v5.0.0) and a qscore cut off of 10 (--min-qscore 10). For high-duplex flow cells, duplex basecalling was run using slow5-dorado (v0.3.4; dna_r10.4.1_e8.2_400bps_sup@v4.2.0) and non-duplex reads were removed prior to downstream analysis. For ONT-only Cornetto experiments, simplex reads were filtered to exclude reads shorter than 30kb (seqkit-v2.3.0)^38^, to avoid excessive coverage within on-target regions (noting that this filtering was not performed on reads used to generate the base assembly).

### Saliva assemblies and nonhuman reads

For human saliva samples, non-human reads were removed before running hifiasm to generate the base assembly. This was performed by running Centrifuge^39^ on the FASTQ input file (see **Supplementary Note 4**). Additionally, non-human contigs were appended to the reference assembly and boringbits during each cornetto iteration, which facilitates rejection of nonhuman DNA during subsequent sequencing. To do this, non-human contigs must be identified by assembling all the reads in the base assembly (both human and non-human) with hifiasm, and then running Centrifuge on the assembly. Any contig in this assembly not assigned with the Homo sapiens species code and covered by a minimum of 100 reads are included (see **Supplementary Note 4**). Centrifuge index used was Bacteria, Archaea, Viruses, Human (compressed) index (p_compressed+h+v).

### Evaluating genome assemblies

To evaluate human genome assemblies, all contigs are first aligned to the Q100 T2T-hg002 (https://github.com/marbl/HG002) paternal haplotype with chrX added, using minimap2 2.24 with preset *asm5* and *--eqx -c* options. Any contigs in the assembly whose sum of aligned lengths for ‘-’ is greater than for ‘+’ with respect to the reference contigs, are reverse complemented. Dot plots are generated using the minidot tool in the miniasm repository^40^. Telomeres are identified in assembled contigs using the telomere analysis script from the VGP project with default options. To obtain per-chromosome metrics of an assembly, each contig in the the assembly is assigned to the corresponding contig in the hg002-paternal reference based on the contig that was most aligned with. Contig L_90_ values are defined as the number of contigs encompassing >90% of the reference sequence for a given chromosome. A chromosome is considered to be complete if it has a L_90_ = 1 and has a telomere detected at both contig ends. All these operations are carried out using wrapper scripts provided in the Cornetto repository (see **Supplementary Note 4**).

To evaluate assembly contiguity, we generated Nx plots by calculating contig sizes and cumulative assembly size for each additional contig, sorted from largest to smallest. N_50_ and N_90_ values are the minimum contig size for which 50% and 90%, respectively, of the genome is assembled into contigs larger than N Mbase. Compleasm 0.2.6^41^ was used for calculating the BUSCO scores with *lineage* set to *primates* for human; *actinopterygii_odb10* for cichlid; and, *tetrapoda_odb10* for both birds and turtles. Yak (v0.1) was used to calculate the QV, hamming error and switch error rates for assemblies. For QV value, separate k-mer count indexes are created using yak count with k-mer sizes 21 and 31 (*-k* option) using the Q100 T2T-hg002 reference genome which includes both paternal and maternal haplotypes. These indexes are used with yak qv subtool to calculate the QV. For calculating hamming and switch errors, first the parental yak indexes for HG002 were downloaded from the human-pangenomics project. Then yak trioeval was used. NGA50, NGA90 value and NGAx plots were obtained using quast 5.2^42^. Assembly guided structural variant calling was performed using dipcall v0.3^43^ against the chm13 reference and evaluation was performed using truvari v5.3.0^44^. Example commands are provided in **Supplementary Note 4**.

During evaluation, primary assemblies the user may optionally refine the assembly by retaining complete chromosomes from earlier cornetto cycles, which are sometimes broken during subsequent cycles. The Cornetto software contains a script to execute this, using the following logic. Suppose the base assembly is called asm-0.fasta and the cornetto iterations are named asm-1.fasta, asm-2.fasta, asm-3.fasta, …, asm-n.fasta. Starting from asm-1.fasta, asm-2.fasta, asm-3.fasta, …, asm-n.fasta are iterated until any contigs are found longer than the expected minimum chromosome size and have telomeres in both ends (considered to be ‘complete chromosomes’). Suppose complete chromosomes are found in asm-k.fasta. Now such contigs are extracted from asm-k.fasta into a file called asm.fasta. Now starting from asm-(k+1).fasta, assemblies are iterated till asm-n.fasta (including asm-n.fasta). At each iteration, any complete chromosomes in the assembly are mapped to asm-k.fasta. Any newly found complete chromosomes are appended to asm.fasta (those contigs which map <50% of their length to a contig into asm.fasta). At the last iteration (asm-n.fasta), any other contigs (that are not considered complete chromosomes) are also mapped to asm.fasta. Any contigs which map <50% of their length to a contig are appended into asm.fasta. At the end, asm.fasta is the final curated primary assembly.

## DATA AVAILABILITY

The raw sequencing data for animals and reference samples generated in this study have been deposited in the European Nucleotide Archive (ENA) under accession code **PRJEB86853**. Human participant data are available under restricted access to protect participant privacy. Participants are enrolled in the Garvan Institute’s Genomics of Rare Disease Registry and enquiries for data access should be addressed to the corresponding author I.W.D. Genome assemblies are available at Dryad (10.5061/dryad.kkwh70sfr). Source data are provided in the **Supplementary Information** (**Supplementary Data 1-7**). The following publicly accessible datasets were also used in this study:

hg002 ONT duplex data from the T2T Consortium:

https://human-pangenomics.s3.amazonaws.com/index.html?prefix=submissions/0CB931D5-AE0C-4187-8BD8-B3A9C9BFDADE--UCSC_HG002_R1041_Duplex_Dorado/Dorado_v0.1.1/stereo_duplex/

hg002 PacBio HiFi data from the T2T Consortium:

https://s3-us-west-2.amazonaws.com/human-pangenomics/T2T/scratch/HG002/sequencing/hifirevio/m84005_220827_014912_s1.hifi_reads.fastq.gz

hg002 HiC data from the HPRC:

https://s3-us-west-2.amazonaws.com/human-pangenomics/working/HPRC_PLUS/HG002/raw_data/hic/downsampled/HG002.HiC_2_NovaSeq_rep1_run2_S1_L001_R1_001.fastq.gz

https://s3-us-west-2.amazonaws.com/human-pangenomics/working/HPRC_PLUS/HG002/raw_data/hic/downsampled/HG002.HiC_2_NovaSeq_rep1_run2_S1_L001_R2_001.fastq.gz

T2TC-ONT-only assembly (Koren et al 50x duplex + 30x UL + PoreC ) assembly:

https://obj.umiacs.umd.edu/marbl_publications/duplex/HG002/asms/duplex_50x_30xUL_poreC.tar.gz

## CODE AVAILABILITY

Cornetto software is open source and freely available: https://github.com/hasindu2008/cornetto

All original code has been deposited at Zenodo and is publicly available: 10.5281/zenodo.15075988.

## ACKNOWLEDGEMENTS

The authors acknowledge the traditional custodians of the land upon which the orange-bellied parrot, western saw-shelled turtle, Gould’s petrel and redstriped eartheater cichlid reside, as well as the custodians of the historic range of each species. We thank Deborah Bower and Yuna Kim for involvement in specimen collection for the turtle and petrel, respectively. We thank Priam Psittaculture Centre for sampling of the parrot. This project was undertaken with services from the National Computational Infrastructure (NCI). We thank Tim Ho for expert technical support.

We acknowledge the following funding support: Australian Medical Research Futures Fund grants, 2023126, 2041648, 2025138, 2008249, National Health and Medical Research Council (NHMRC) grant 2035037 (to I.W.D.), Australian Research Council (ARC) DECRA Fellowship DE230100178 and ARC Discovery Project DP230100651 (to H.G.). A.J.M was supported by a Queensland Health Advancing Clinical Research Fellowship. L.M.S. is supported by the ARC Centre of Excellence in Innovations in Peptide and Protein Science (CE200100012). R.C.R.N and L.A.S.N were supported by the National Council for Scientific and Technological Development (grants 444448/2024-0, 306665/2025-5 and 309711/2023-1) and Amazonia Foundation for Support of Studies and Research -FAPESPA (grant 167/2024). D.Y. is supported by a PhD Scholarship from Muscular Dystrophy NSW. K.R.K. is supported by the Ainsworth 4 Dystonia Genetic Research Mission. Work on the orange-bellied parrot was supported by Australian Biocommons which is enabled by National Collaborative Research Infrastructure Scheme via Bioplatforms Australia Threatened Species Initiative funding; DNRF143. The views expressed herein are those of the authors and are not necessarily those of the Australian Government or the ARC, NHMRC or MRFF.

## AUTHOR CONTRIBUTIONS STATEMENT

H.G., H.R.P. & I.W.D. conceived the project. H.G., K.J., & I.W.D. developed the Cornetto software. I.S., J.M.H., M.R., T.R. & Y.H. performed laboratory experiments. I.S., D.Y., Y.H., A.J.M., E.S., C.P., K.R.K. & A.C.M. recruited, collected and processed patient samples. J.M.H., L.R., L.W.S., C.J.H., L.M.S., O.B., R.C.R.N., L.A.S.N., A.L.C. & A.G. coordinated, collected and processed non-human samples. H.G., I.S., J.M.H., A.L.M.R., L.W.S., D.Y., H.C., H.R.P. & I.W.D. performed bioinformatics analysis. H.G., A.L.M.R., D.Y., A.C.M. & I.W.D. prepared the figures and tables. H.G., I.S. & I.W.D. wrote the manuscript, with contributions from all co-authors.

