## Supplementary Materials for "Targeted sequencing and iterative assembly of near-complete genomes"

### SUPPLEMENTARY INFORMATION

**Supplementary Data 1.** Comparison of hg002 assemblies from PacBio HiFi and ONT duplex data. *[Excel file]*

**Supplementary Data 2.** Comparison of primary assemblies for hg002 from ONT data. *[Excel file]*

**Supplementary Data 3.** Comparison of diploid assemblies for hg002 from ONT data. *[Excel file]*

**Supplementary Data 4.** Assembly-guided structural variant detection. *[Excel file]*

**Supplementary Data 5.** Comparison of genome assemblies from human saliva. *[Excel file]*

**Supplementary Data 6.** Comparison of genome assemblies for non-human vertebrates. *[Excel file]*

**Supplementary Data 7.** Compute resources used during genome assembly. *[Excel file]*

**Supplementary Figure 1.** Comparison of long-read sequencing data types used for assemblies. page 1

**Supplementary Figure 2.** ONT-only human genome primary assemblies. page 2

**Supplementary Figure 3.** Diploid human genome assemblies for hg002. page 3

**Supplementary Figure 4.** Saliva assemblies generated with HiFi and duplex data. page 4

**Supplementary Figure 5.** Genome assemblies for non-human vertebrates. page 5

**Supplementary Note 1**. Improving genetic diagnosis of FSHD page 6-8

**Supplementary Note 2**. Improving genetic diagnosis of *MUC1*-ADTKD page 9-12

**Supplementary Note 3**. Genome assemblies for selected non-human vertebrates page 13-15

**Supplementary Note 4**. Extended methods page 16-19

**Supplementary References** page 20

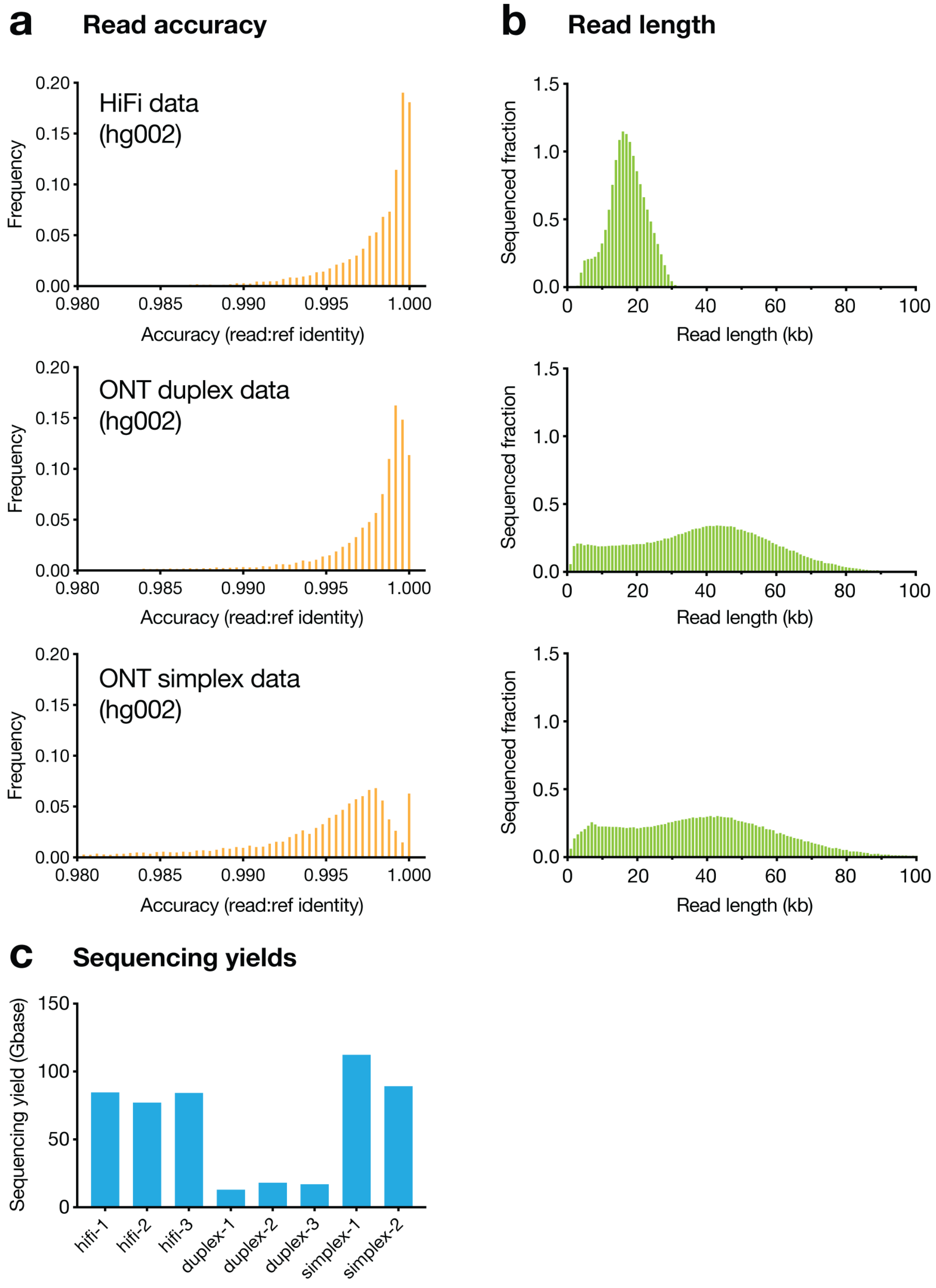


**Supplementary Figure 1. Comparison of long-read sequencing data types used for assemblies.** Plots compare relevant metrics for PacBio HiFi, ONT duplex and ONT simplex datasets generated with hg002 DNA and used to generate genome assemblies described throughout. (**a**) Histograms show read-level sequencing accuracy, assessed based on read:reference identity scores for alignments to the Q100 T2T-hg002 reference. (**b**) Histograms show read length distributions, expressed as the fraction of total sequencing data represented in each sequential 1kb read-length bin. (**c**) Bar chart shows sequencing yields obtained with individual PacBio SMRT cells, ONT duplex flow cells and ONT simplex flow cells. Yields are the cumulative length of all reads meeting HiFi quality for PacBio and ONT pass reads, and are shown for ONT flow cells run without Cornetto.

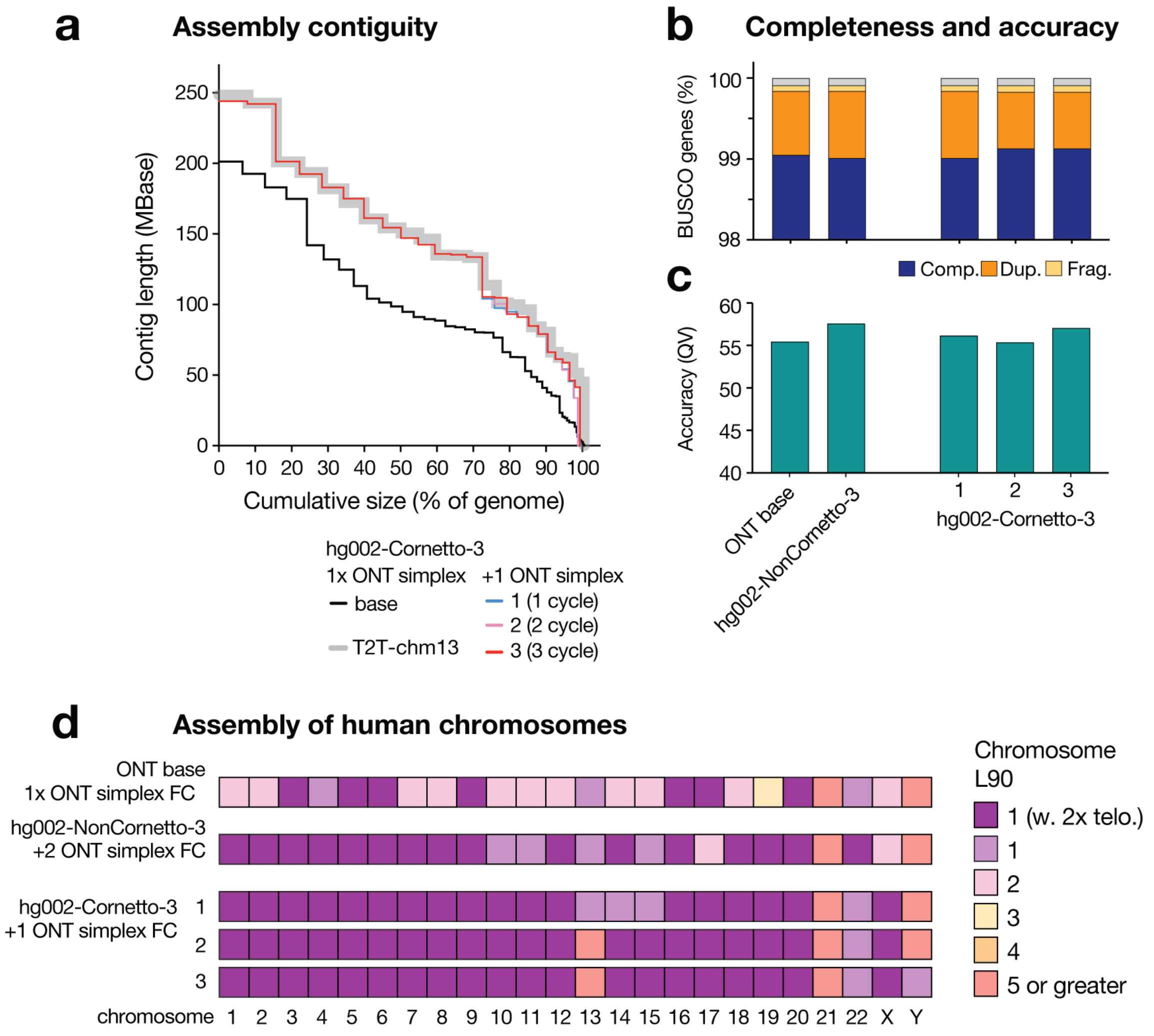


**Supplementary Figure 2. ONT-only human genome primary assemblies**. (**a**) Nx plots show contigs sizes sorted from largest to smallest, relative to cumulative assembly size, as a percentage of the human genome size (3.1 Gbase). T2T-chm13 genome is shown for comparison (grey line). Assemblies shown are a base assembly generated using one standard ONT flow cell (LSK114; simplex data) compared to the *hg002-Cornetto-3* assembly, generated by running the same library on one additional ONT flow cell. Three separate primary assemblies are shown for *hg002-Cornetto-3*, which represent the intermediate assemblies generated at each Cornetto pause-point. (**b**) Stacked bar charts show the proportion of BUSCO genes detected as complete, duplicated, fragmented or missing for *hg002-Cornetto-3* (three intermediate assemblies) compared to *hg002-NonCornetto-3*, which was generated using 3x ONT flow cells in total. (**c**) Bar chart shows assembly sequence accuracy, as per QV values (k-mer size of 21). (**d**) Tile plot shows contiguity of human chromosomes in same assemblies. Colour scale encodes $L_{90}$ values: number of contigs encompassing >90% of the reference sequence for a given chromosome. Dark purple tiles show chromosomes with $L_{90} = 1$ and a telomere detected at each end, indicating the whole chromosome is assembled as a single primary contig.

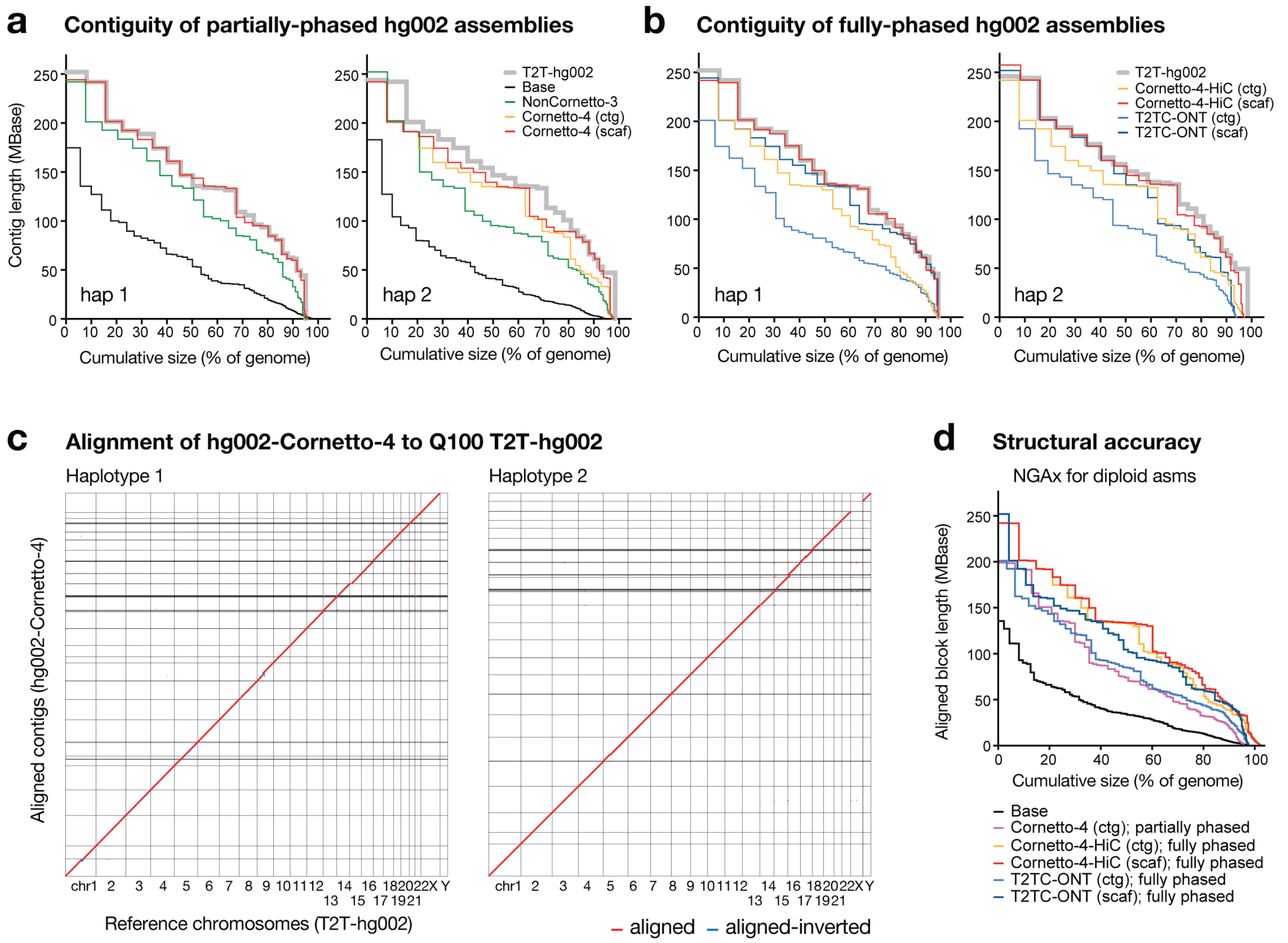


| | | |
|---|---|---|
| hg002 assemblies | Base = ONT simplex (1x FC) | NonCornetto-3 = ONT simplex (3x FC) |
| | Cornetto-4 = ONT simplex (2x FC) | Cornetto-4-HiC = ONT simplex (2x FC) + Hi-C (1x FC) |
| | T2TC-ONT = ONT duplex (10x FC) + ultra-long (2x FC) + pore-C (3x FC) | |

**Supplementary Figure 3. Diploid human genome assemblies for hg002.** (**a**,**b**) Nx plot shows contig/scaffold sizes sorted from largest to smallest, relative to cumulative assembly size, as a percentage of the human genome size (3.1 Gbase). The plot compares the diploid assemblies generated for hg002 with haplotypes plotted separately. Partially-phased (**a**) and fully-phased (**b**) assemblies are plotted separately and scaffolds vs contigs are shown where relevant. Data types and sequencing resources used for each assembly are detailed in the legend at the bottom. A published ONT-only assembly from the T2T Consortium is included for comparison (T2TC-ONT). The Q100 T2T-hg002 assembly provides a reference. (**c**) Dot plot shows the alignment of contigs in the diploid *hg002-NonCornetto-4* assembly (vertical axis) to chromosomes in Q100 T2T-hg002 (horizontal axis). Each *hg002-NonCornetto-4* haplotype was aligned separately to the appropriate T2T-hg002 haplotype. (**d**) NGAx plot shows the length of contig/scaffold alignments to the T2T-hg002 reference, relative to cumulative assembly size. Both haplotypes are included during the alignment and cumulative size is shown as a percentage of the diploid human genome size (6.2 Gbase). Assemblies included are the same as for **a** and **b**, with details listed in the legend below.

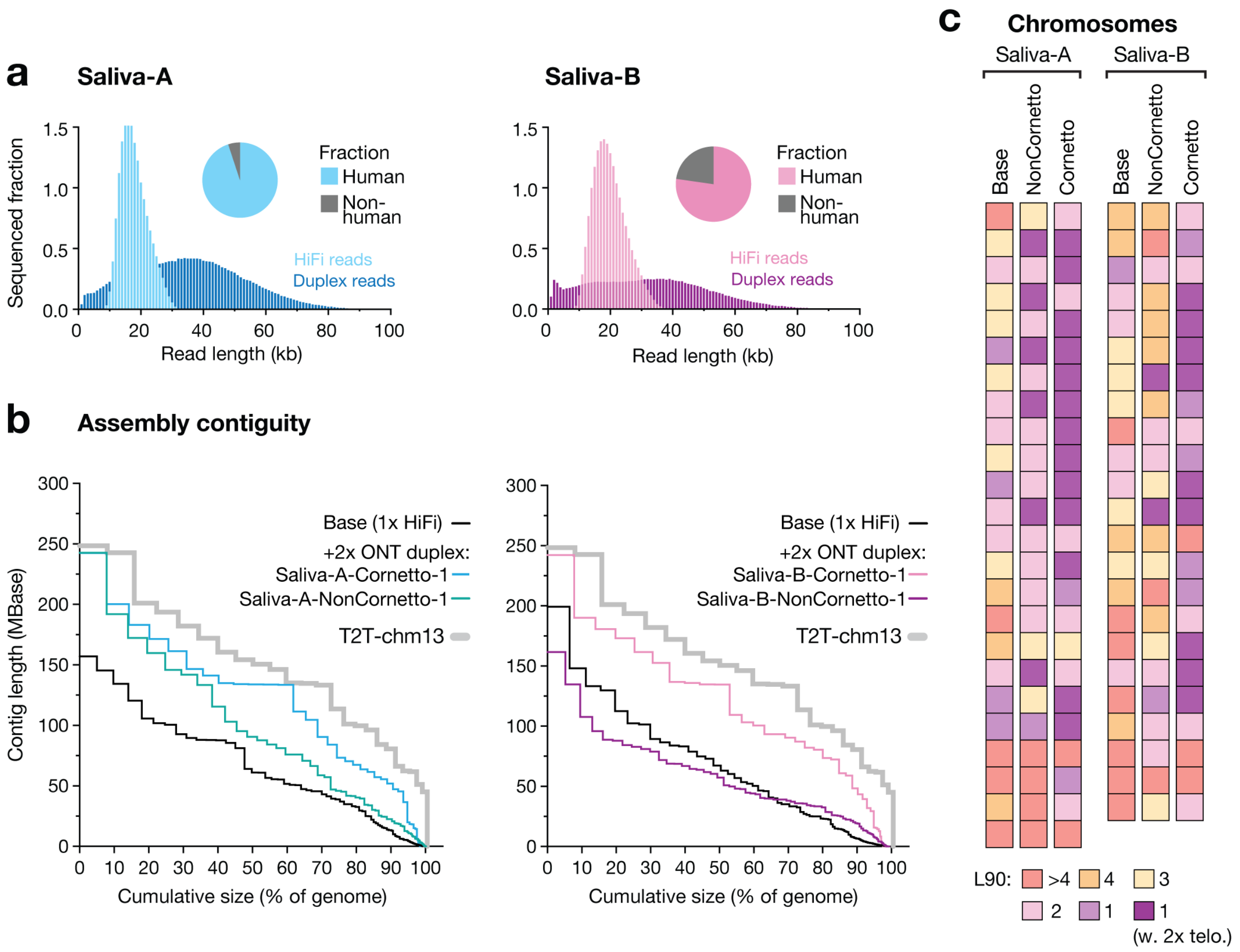


**Supplementary Figure 4. Saliva assemblies generated with HiFi and duplex data.** (**a**) Histograms show read length profiles for PacBio HiFi data and ONT duplex data on saliva samples from two participants: Saliva-A (male) and Saliva-B (female). Pie charts show the proportion of non-human reads in the HiFi dataset used to generate the base assembly for each individual. (**b**) For a given primary assembly, Nx plots show contigs sizes sorted from largest to smallest, relative to cumulative assembly size, as a percentage of the human genome size (3.1 Gbase). For each participant, assemblies shown are the base assembly generated with PacBio HiFi data from 1 SMRT cell; a Cornetto assembly generated using 1 SMRT Cell and 2 duplex flow cells (Saliva-A-Cornetto-1 / Saliva-B-Cornetto-1; 3 cycles per flow cell); a comparative assembly generated using 1 SMRT cell and 2 duplex flow cells run without Cornetto (Saliva-A-NonCornetto-1 / Saliva-B-NonCornetto-1). T2T-chm13 reference is shown for comparison on each plot. (**c**) Tile plot shows contiguity of human chromosomes in same assemblies. Colour scale encodes $L_{90}$ values: number of contigs encompassing >90% of the reference sequence for a given chromosome. Dark purple tiles show chromosomes with $L_{90}$ = 1 and a telomere detected at each contig end, indicating the whole chromosome is assembled as a single primary contig.

Genome assemblies for non-human vertebrates

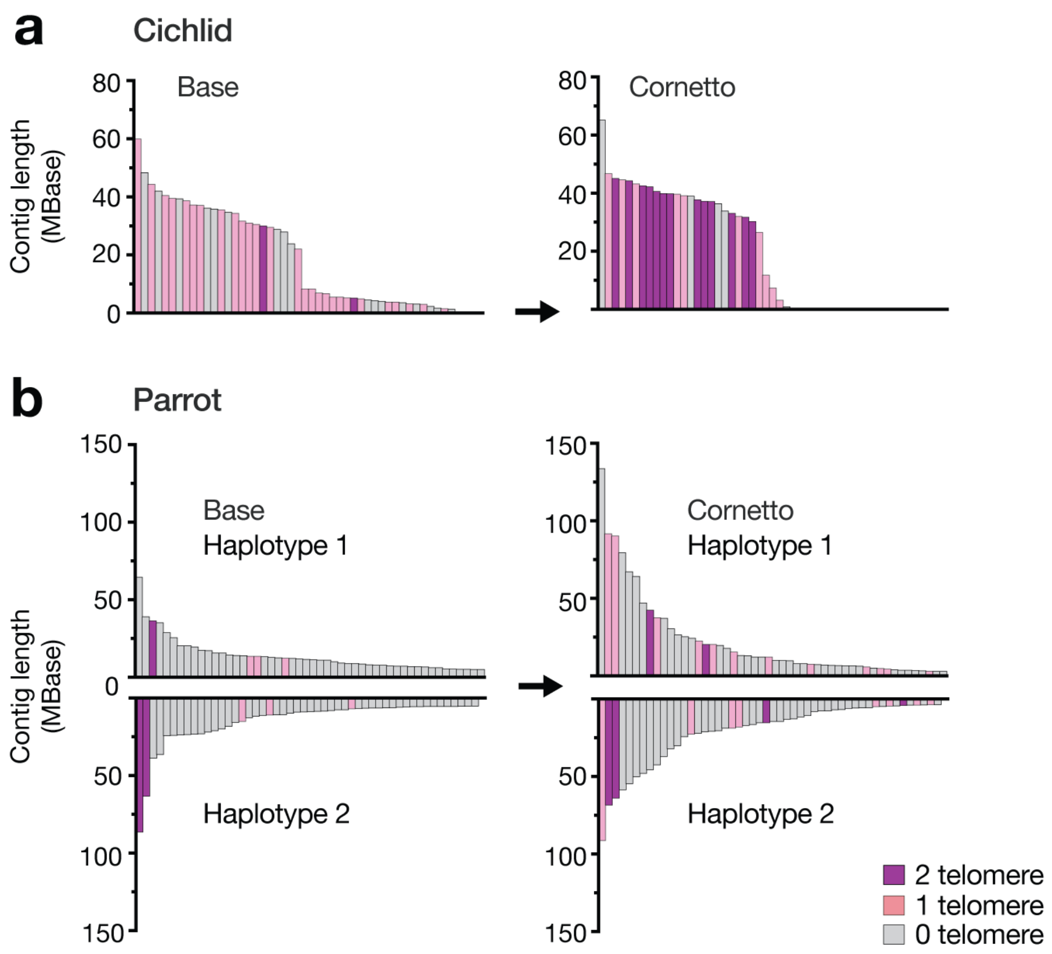


**Supplementary Figure 5. Genome assemblies for non-human vertebrates.** (**a,b**) Bar plots show sizes of the fifty largest contigs in descending order, coloured according to presence of telomere sequences at contigs ends (both ends = purple; one end = pink). (**a**) Primary assemblies are shown for the cichlid, which was assembled with PacBio HiFi (base; 1 SMRT cell) plus ONT duplex data (cornetto; 2 duplex cells). (**b**) Diploid assemblies are shown for the orange-bellied parrot, which was assembled using ONT data, two standard flow cells (base) plus a second with Cornetto (simplex reads; LSK114). Haplotypes are shown separately for the diploid assembly. Equivalent plots for the petrel and turtle are shown in **Figure 6**.

#### SUPPLEMENTARY NOTE 1 – Improving genetic diagnosis of FSHD

Facioscapulohumeral muscular dystrophy (FSHD) is a progressive myopathy caused by aberrant expression of the *DUX4* gene buried within the D4Z4 macrosatellite repeat located in the subtelomeric region of chromosome 4q. The repeat typically ranges in size from ~11–100 copies[1]. Sequence variation distal to D4Z4 designate each 4q allele as being of one of two major haplotypes: 4qA or 4qB. The most common form of FSHD, FSHD type 1 (FSHD1), is caused by a contraction in the D4Z4 repeat (~1-10 copies) that must be *in cis* with a permissive 4qA allele – this facilitates expression of the *DUX4* gene[1]. FSHD type 2 (FSHD2) is a less common, digenically inherited form of FSHD that is clinically indistinguishable from FSHD1. FSHD2 is caused by standard pathogenic variants in one of three genes involved in DNA methylation (*SMCHD1*, *DNMT3B*, or *LRIF1*) in a patient who has at least one 4qA allele[1]. The 4qA D4Z4 repeat size in FSHD2 is typically in the lower normal range (up to ~20 D4Z4 repeats).

Currently, a genetic diagnosis of FSHD1 is most frequently obtained by performing a Southern Blot assay, in which DNA is digested by specific restriction enzymes and separated by gel electrophoresis, followed by estimation of D4Z4 repeat number based on size of the Southern Blot fragment. Determination of 4q haplotype is performed using different restriction enzymes and specific 4qA or 4qB probes. The large size and repetitive nature of the D4Z4 repeat array and the presence of a highly homologous D4Z4 repeat array on chromosome 10 means that this locus is refractory to characterisation by short-read sequencing. Furthermore, Southern Blot analysis may be negative or inconclusive in cases of FSHD2 or in the presence of more complicated genetic variants e.g. somatic mosaicism for D4Z4 repeat contraction, structural variants (e.g. *in cis* duplications) of the D4Z4 repeat array, proximal extended D4Z4 deletions (that remove Southern blot probe binding sites) and translocations between the D4Z4 arrays on chromosome 4 and 10.

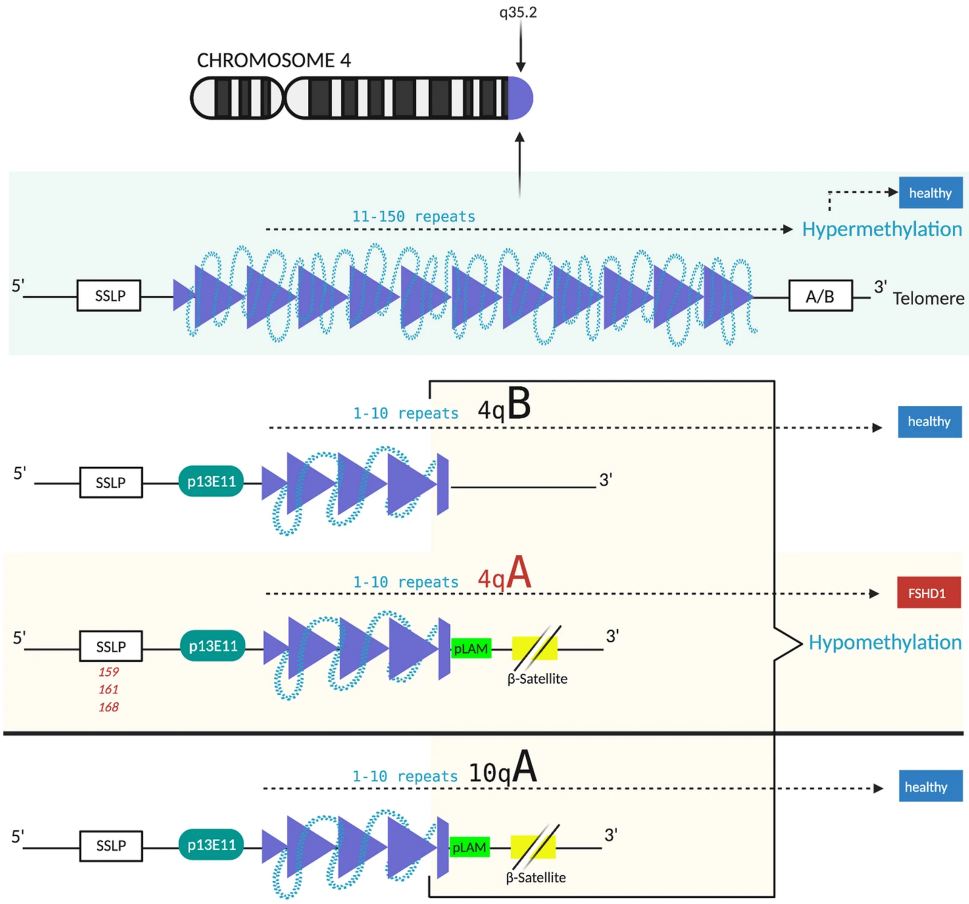


Schematic from Schätzl, Kaiser & Deigner (2021)[2] summarising the genetic basis for FSHD. >10 D4Z4 repeats leads to a high methylation of D4Z4 macrosatellite and represses transcription of DUX4. There are two possible D4Z4 haplotypes A and B, which are equally common. FSHD only occurs in individuals that carry the permissive 4qA allele. FSHD manifests in people with D4Z4 contraction and carrying 4qA, whereas 4qB and are healthy. Haplotype A can be identified by the pLAM sequence.

To assess the capacity of Cornetto to resolve D4Z4, we retrieved the 4q subtelomeric region from both haplotypes in the hg002-Cornetto-4 assembly. We genotyped the locus by mapping of known sequence features relevant to FSHD, including canonical D4Z4 repeat elements, pLAM, SSLP, p13E11 and sequence markers for 4qA vs 4qB haplotypes[2]. We identified one 4qA haplotype with 42 copies and one 4qB haplotype with 26 copies. Comparison of our *hg002-Cornetto-4* assembly to the *T2T-hg002* Q100 reference genome showed perfect concordance in the size and near-perfect sequence identity between 4q D4Z4 haplotypes, indicating these are assembled with very high accuracy (see **Fig3a** in the main manuscript). Therefore, targeted ONT sequencing and haplotype-aware assembly of the *D4Z4* locus may be a viable strategy for the genetic diagnosis of FSHD.

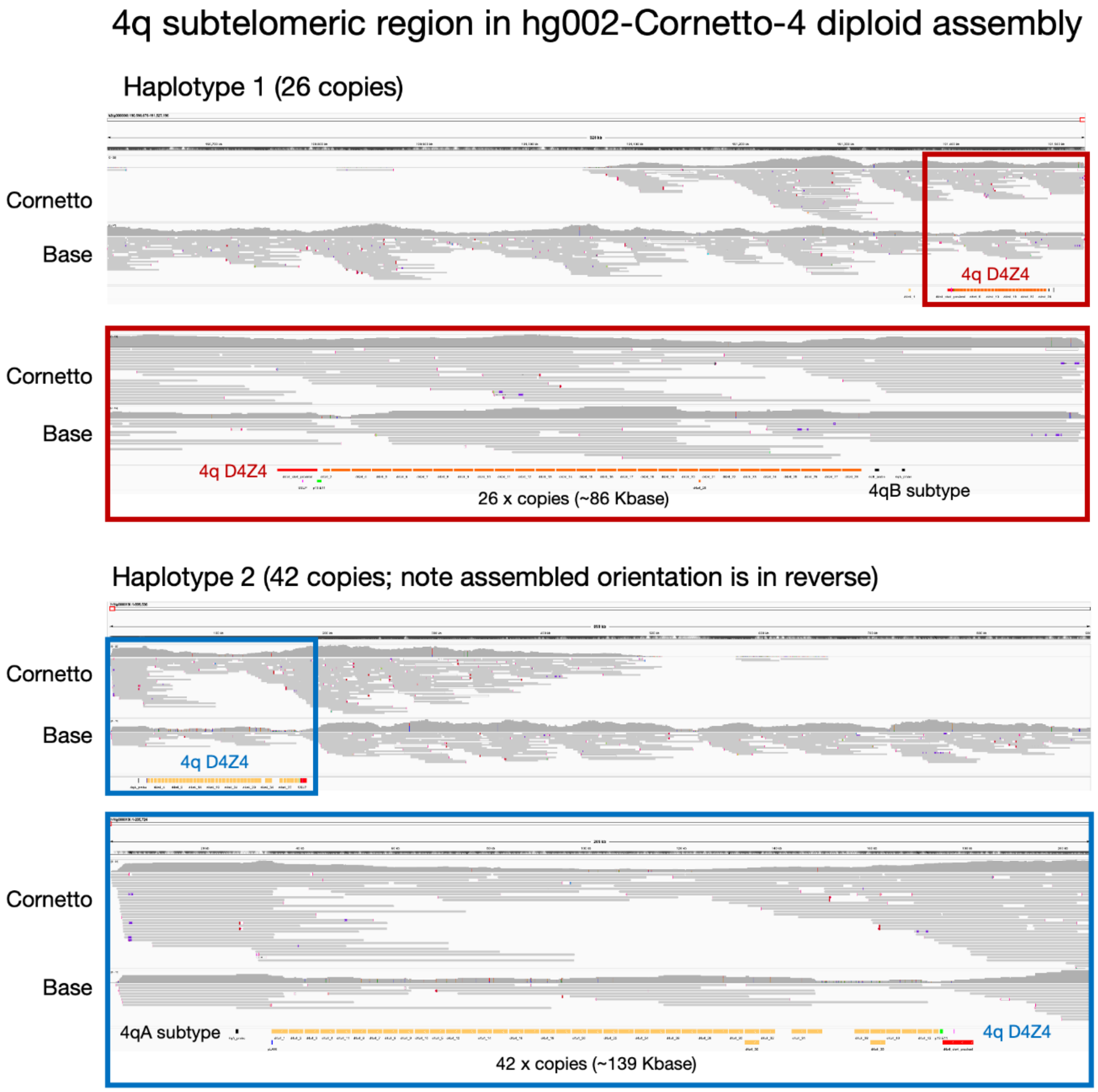


Browser views show analysis of the D4Z4 locus in the *hg002-Cornetto-4* diploid assembly. ONT reads from the base library and Cornetto iterative sequencing are re-aligned to the assembly, demonstrating good coverage with long, uniquely aligned reads to each D4Z4 haplotype. Mapping of known sequence features including D4Z4 repeat elements, pLAM, SSLP, p13E11 and sequence markers for 4qA vs 4qB haplotypes allows us to genotype the locus, revealing one 4qA haplotype with 42 copies (lower) and one 4qB haplotype with 26 copies (upper). A close-up view of the detected pLAM sequence, identifying the 4qA haplotype is shown over the page.

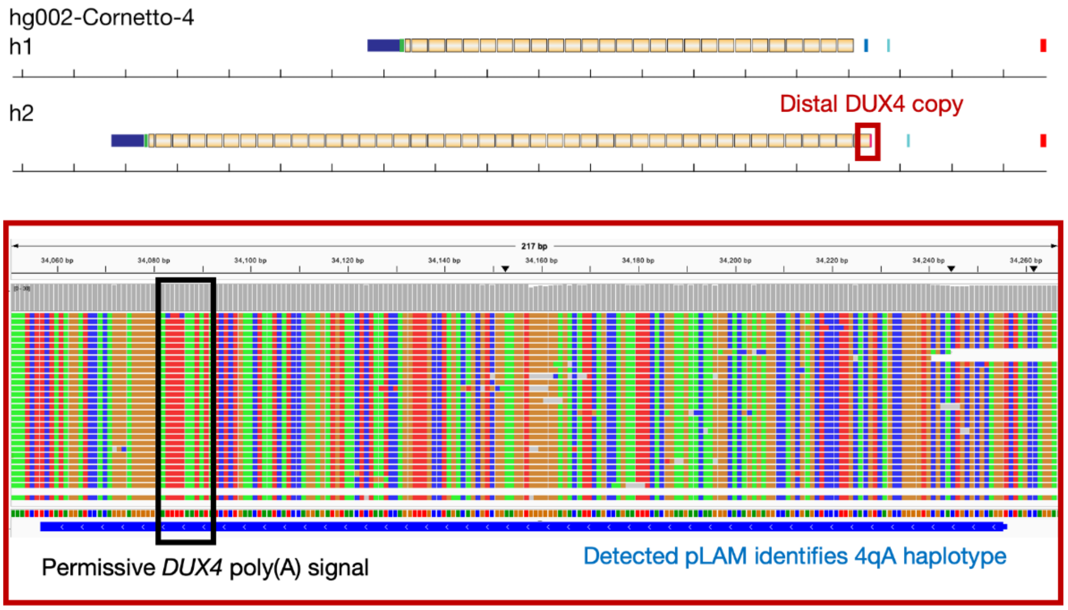


Browser view shows accurate detection of the permissive 4qA allele at the distal end of the D4Z4 repeat. 4qA is defined by the pLAM sequence, which carries a poly-adenylation signal required for *DUX4* expression. Alignments to the second haplotype in the hg002-Cornetto-4 assembly show good coverage and strong sequence consensus across the pLAM.

To test this further, we created a target panel for programmed targeted sequencing and genotyping of the *D4Z4* locus. This is a more streamlined approach than enrichment and assembly of unsolved genome regions globally, as per the standard Cornetto paradigm, and would hence be preferable in a diagnostic context. However, we note the target regions are a subset of regions enriched during standard Cornetto sequencing.

To validate this method, we performed targeted ONT sequencing and *D4Z4* assembly for four patients with a known diagnosis of FSHD1. Prior testing consisted of pulsed-field gel electrophoresis-Southern blot showing pathogenic repeat contractions (4, 5, 6 and 9 repeats). Two patients also had 4q haplotyping confirming the contracted repeat was on a permissive 4qA allele. Clinical features for these patients, information from previous genetic testing, and results from targeted ONT assembly are summarised in the table below. In each case we identified one D4Z4 haplotype of the permissive 4qA sub-type with fewer than 10 copies, sufficient for a positive diagnosis. We observed 4q D4Z4 lengths that were concordant with previous molecular genetic testing (see **Fig3a** in the main manuscript). These results indicate targeted assembly of the D4Z4 region is a viable strategy for genetic diagnosis of FSHD.

**Clinical features and genetic testing results of FSHD1 patients analysed by ONT sequencing**

| Patient | Clinical features | FSHD PFGE-SB Fragment Size (Estimated D4Z4 Repeat Number) and 4q Haplotype* | D4Z4 Repeat Number and 4q Haplotype Derived from Targeted ONT Sequencing |
|---|---|---|---|
| FSHD p1 | Mild facial weakness, asymmetric scapular winging and shoulder girdle muscle weakness | 33 kb (9 repeats) on a 4qA allele | 9 D4Z4 copies (29.7 kb) on a 4qA allele |
| FSHD p2 | Asymmetric scapular winging and shoulder girdle muscle weakness | 23 kb (6 repeats); 4q haplotype not tested | 6 D4Z4 copies (19.8 kb) on a 4qA allele |
| FSHD p3 | Asymmetric facial weakness, scapular winging, shoulder girdle muscle and ankle dorsiflexor weakness | 19 kb (5 repeats); 4q haplotype not tested | 5 D4Z4 copies (16.5 kb) on a 4qA allele |
| FSHD p4 | Severe facial diplegia, bilateral scapular winging, bilateral shoulder girdle, hip flexor and ankle dorsiflexor muscle weakness | 17 kb (4 repeats) on a 4qA allele | 4 D4Z4 copies (13.2 kb) on a 4qA allele |

FSHD, facioscapulohumeral muscular dystrophy; ONT, Oxford nanopore technology PFGE-SB, pulsed-field gel electrophoresis-Southern blot | *fragment size determined by *EcoRI*/*BlnI* double digest PFGE-SB using p13E-11 probe; presence of 4qA determined by *HindIII* digest PFGE-SB using 4qA-specific probe.

#### SUPPLEMENTARY NOTE 2 – Improving genetic diagnosis of *MUC1*-ADTKD

Autosomal Dominant Tubulointerstitial Kidney Disease (ADTKD) is predominantly caused by pathogenic variants in one of four genes – *UMOD, MUC1, REN* and *HNF1B* – and is among the most common monogenic forms of renal disease[3,4]. The condition is characterised by bland urinary sediment, fibrosis of the kidney tubulointerstitial tissue and chronic kidney disease, with kidney failure reported throughout adulthood[5,6]. Disease due to pathogenic variants in the *MUC1* gene (*MUC1*-ADTKD) are thought to account for approximately 20% of ADTKD, though the true prevalence of *MUC1*-ADTKD is unknown due to significant technical challenges in diagnosing the condition[3].

The *MUC1* gene contains a complex variable number tandem repeat region (VNTR)[7]. The *MUC1* VNTR is composed of recurring 60 bp imperfect sequence subunits, with the number of subunit copies being highly variable between individuals (20-125 units per allele)[7,8]. In addition, the composition of sequence subunits differs within and between individuals[7,8]. A frameshift variant (dupC) resulting from duplication of a cytosine on a 7xC homopolymer in the wild-type sequence has been identified as the most common mechanism of disease in MUC1-ADTKD[6], with a small handful of other rare variants also reported[4]. Differences in the length of the VNTR are not disease-causing and not known to be correlated with the severity or nature of symptoms.

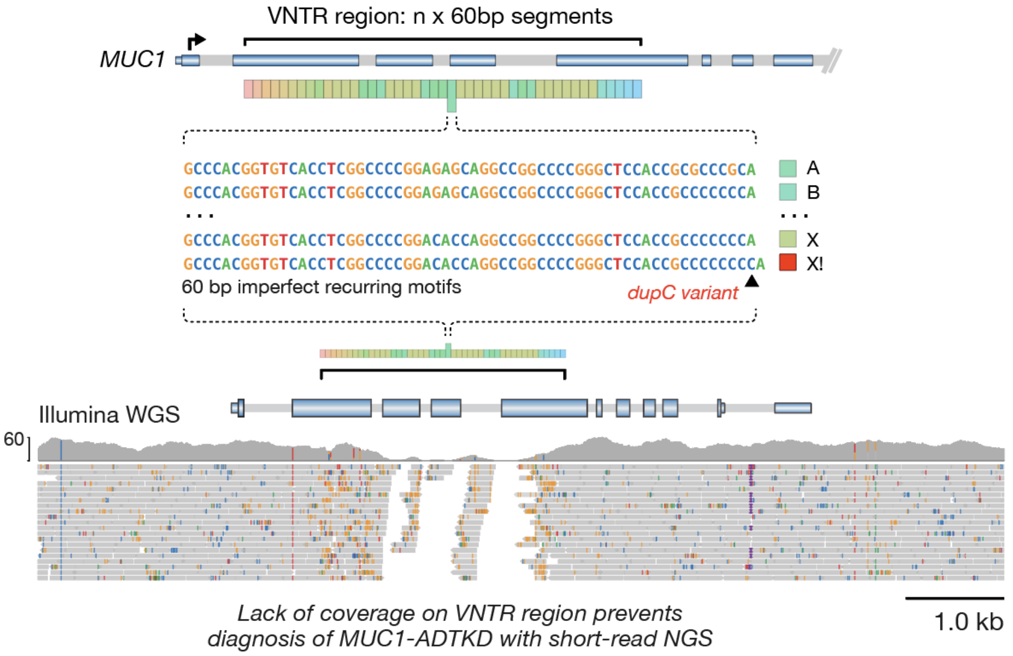


Schematic diagram showing architecture of MUC1 VNTR region and 'dupC' variant. Genome browser view underneath shows inability to resolve MUC1 VNTR with short-read sequencing.

Due to the complexity of the VNTR sequence, genetic diagnosis has been challenging in *MUC1*-ADTKD, highlighted by the genetic basis of the disease only being identified in 2013. A probe extension assay combined with mass spectrometry has been the mainstay of diagnosis[7]. This assay is technically intensive, making availability of diagnostic testing largely limited to a single laboratory, worldwide. The highly repetitive nature and size of the VNTR, along with high GC content (>80%), confounds analysis by short-read next-generation sequencing (NGS). Recently, new bioinformatic tools have had some success in identifying specific cytosine duplications within the VNTR using short-read NGS data[9,10,11]. However, these are limited to detect specific sequence variants and cannot place these within a complete VNTR sequence, nor identify which haplotype harbours the variant. PacBio LRS (using HiFi or previous generation circular consensus sequencing [CCS]) has been trialled, though due to cost this has been performed from PCR-amplified DNA, increasing laboratory complexity and potential for introduction of PCR-errors[12].

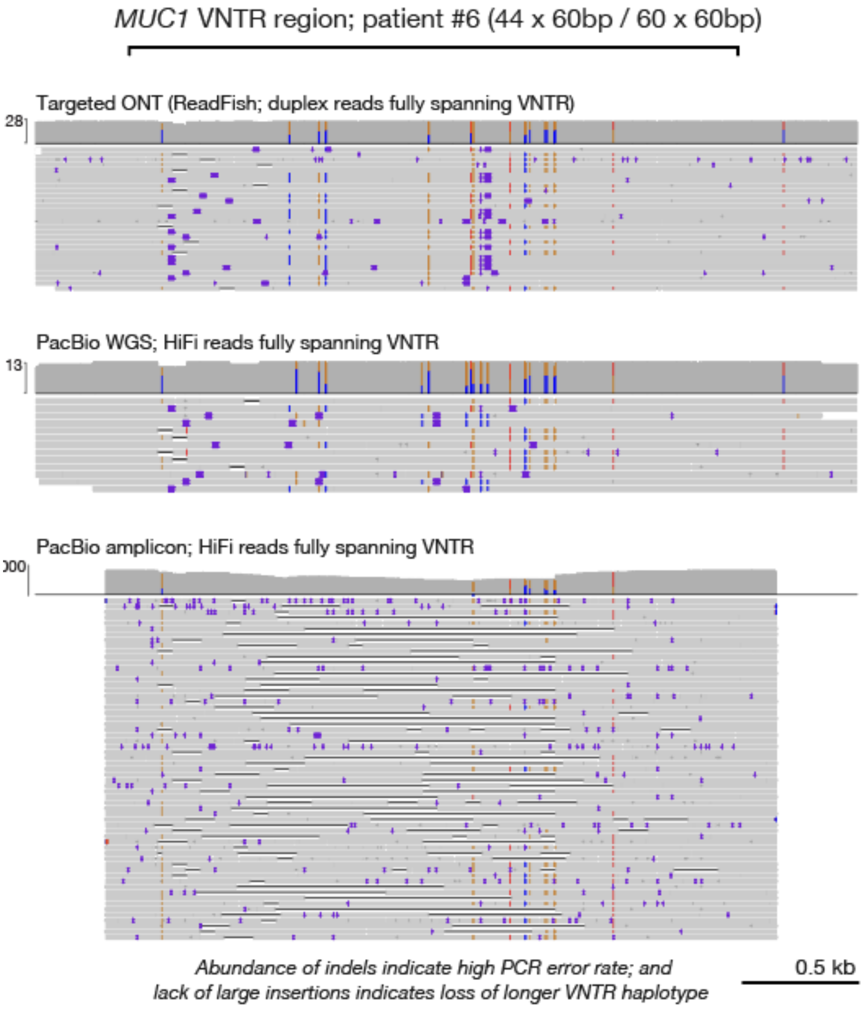


Browser views show alignment of different long-read sequencing data types to the MUC1 VNTR region. All data is from a single patient (patient 6), who harbours VNTR haplotypes of 44 x 60bp and 60 x 60bp length. Only alignments fully spanning the VNTR sequence are shown. Top track shows ONT duplex reads enriched using programmable selective sequencing (ReadFish or Adaptive Sampling). Data from one flow cell is shown. Middle track shows whole-genome PacBio HiFi sequencing. Combined data from 2x SMRT cells is shown. Bottom track shows PacBio HiFi data enriched by PCR amplification of the VNTR region. While PCR amplification generates high coverage of the locus, the data is characterised by a very high rate of PCR errors, manifesting as both small and large indels. Additionally, the absence of large insertions that are visible in the two upper tracks (which are amplification-free) indicates the larger VNTR haplotype is lost during PCR amplification.

Developing a reliable diagnostic method is crucial for *MUC1*-ADTKD. The true prevalence is unknown due to markedly limited international diagnostic options for this disease. This results in patients being undiagnosed and limits affected families' access to early diagnosis and supportive treatments, along with opportunity to use genetic diagnosis to inform their reproductive planning. Importantly, it has recently been identified in animal and organoid models, that the fibrosis-inducing protein accumulated in patients with a frameshifting variant in MUC1 can be degraded by a novel compound[13]. There is no current treatment for *MUC1*-ADTKD. These findings mean clinical trials are a very realistic short-term possibility in this condition and further reiterates the need for early and readily-available genetic diagnosis worldwide.

Comparison of our diploid *hg002-Cornetto-4* assembly to the *T2T-hg002* Q100 reference genome showed perfect concordance in size and sequence of the MUC1 VNTR haplotypes, indicating these are assembled with very high accuracy (see **Fig4b** in the main manuscript). Therefore, targeted ONT sequencing and haplotype-aware assembly of the *MUC1* locus may be a viable strategy for genetic diagnosis of *MUC1*-ADTKD.

To test this further, we created a target panel for programmed enrichment of the *MUC1* locus and surrounding regions and a custom *MUC1* assembly method. This is a more streamlined approach than enrichment and assembly of unsolved genome regions globally, as per the standard Cornetto paradigm, and would hence be preferable in a diagnostic context. However, we note the target regions are a subset of regions enriched during standard Cornetto sequencing. Our custom assembly strategy is as follows: align reads to T2T-chm13 reference genome; retrieve all alignments that fully span the *MUC1* VNTR region; determine the length of the *MUC1* VNTR sequence within each full-spanning read; organise reads into two groups based on their observed VNTR lengths, representing the two haplotypes for that individual (which are distinct in all cases we have observed so far); align each group of reads to a reference sequence for the *MUC1* VNTR (extracted from T2T-chm13) and run consensus polishing with the Racon package (https://github.com/isovic/racon), producing a polished consensus VNTR sequence for each haplotype, which can then be annotated by aligning previously common 60bp sequence subunits to the consensus sequence, including dupC.

To validate this method, we performed targeted ONT duplex sequencing and *MUC1* assembly in ten patients with diagnostically confirmed *MUC1*-ADTKD. Clinical features for these patients, information from previous genetic testing, and results from targeted ONT assembly is summarised in the table below. Across 11 individuals (including hg002), we observed 20 unique VNTR haplotypes which ranged in size from 40–83 copies, with no individuals sharing the same pair of haplotypes (see **Fig4b** in the main manuscript). In each patient we identified a single dupC frameshift variant within the VNTR occurring on a single haplotype, sufficient for a positive diagnosis. For further validation we performed deep (~50-60X; 2x SMRT cells) PacBio HiFi whole-genome sequencing data on three patients. For each patient, the dupC variant detected by cornetto was supported by HiFi data and identical haplotypes were assembled. It should be noted that, due to the challenging nature of MUC1 (>80% GC content; highly repetitive), we obtained low depth of HiFi reads spanning the locus, by comparison to ONT duplex sequencing with Cornetto.

In summary, these results establish the capacity to accurately assemble the *MUC1* locus in a haplotype-specific fashion and detect the dupC variant, thereby providing a viable new strategy for genetic diagnosis of *MUC1*-ADTKD.

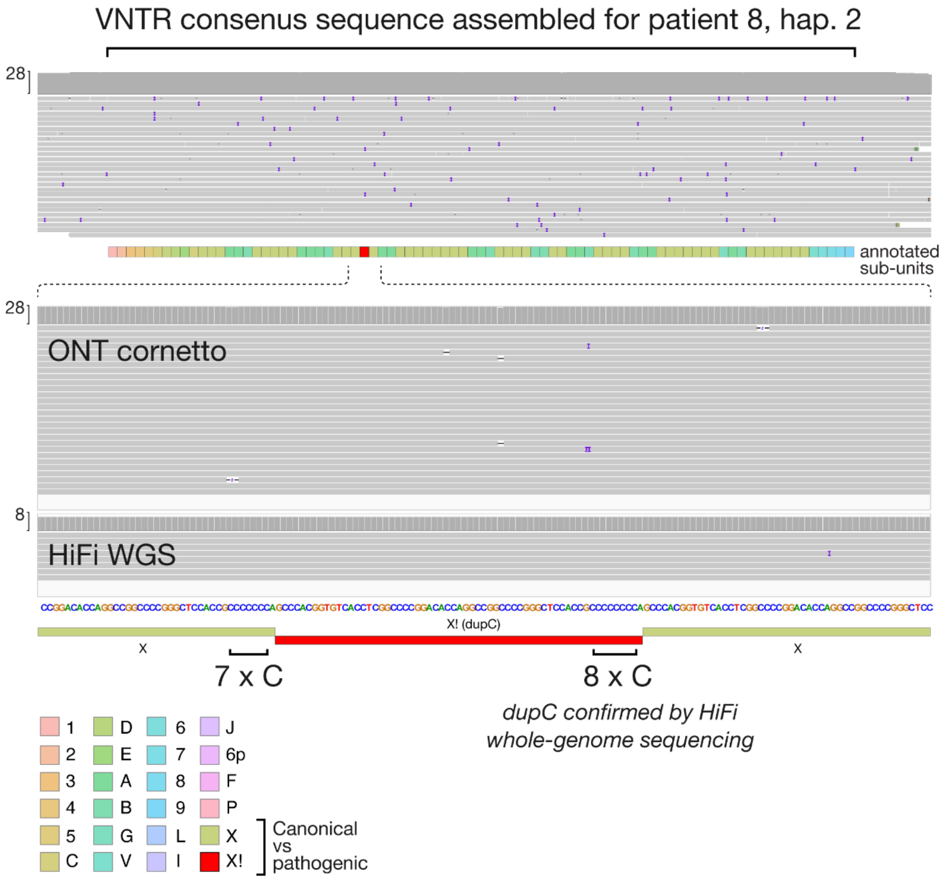


Browser view shows alignment of constituent ONT data to the assembled consensus sequence for one patient haplotype (patient 8, haplotype 2) in which the pathogenic dupC variant was detected. Upper panel shows annotated sequence subunits across the length of the assembled VNTR. Lower panel shows close-up view of the subunit containing the dupC variant (X!) and neighboring canonical (X) subunits. Deep coverage (28x) and strong sequence concordance between alignments provides strong evidence for detection of the dupC variant. In the lower panel, PacBio HiFi whole-genome sequencing data is also shown, which confirms the presence of the dupC variant.

##### Clinical features and genetic investigation in MUC1 cohort

| Patient | Age at recruit. | CKD Stage / Age at kid. failure | Family History Summary | Phenotype Summary | Previous Genetic Investigation | Assembled MUC1 haplotypes |
|---|---|---|---|---|---|---|
| MUC1 p1 | 47 | kidney failure / 31yo | multigeneration family history of CKD | Presented with CKD at 19yo; kidney failure at 31yo; bland urinary sediment; tubulointerstitial disease on biopsy | MUC1-ADTKD diagnosed via Probe extension Assay* | 56 copies (1x dupC) 72 copies normal |
| MUC1 p2 | 49 | | multigeneration family history of CKD | | MUC1-ADTKD diagnosed via Probe extension Assay* | 78 copies (1x dupC) 83 copies normal |
| MUC1 p3 | 54 | kidney failure / 45yo | multigeneration family history of CKD | Presented with CKD at 43yo | MUC1-ADTKD diagnosed via Probe extension Assay* | 44 copies (1x dupC) 46 copies normal |
| MUC1 p4 | 39 | CKD III at 39yo | multigeneration family history of CKD | Diagnosed with CKD at 16yo; bland urinary sediment | MUC1-ADTKD diagnosed via Probe extension Assay* | 66 copies normal 83 copies (1x dupC) |
| MUC1 p5 | 75 | ESKD / 33yo | multigeneration family history of CKD | Diagnosed with CKD at 32yo | MUC1-ADTKD diagnosed via Probe extension Assay* | 44 copies normal 82 copies (1x dupC) |
| MUC1 p6 | 33 | kidney failure / 31yo | unaffected mother and father (molecularly confirmed as de novo) | CKD III at 28yo; bland urinary sediment; renal biopsy at presentation - moderate to severe interstitial fibrosis and tubular atrophy; ultrasound imaging at presentation - 10.9cm right kidney length and 10cm left kidney length with some simple renal cysts; abdominal MRI at presentation - right kidney - 7 cysts, largest 8mm, left kidney 6 subcentimeter cysts; | MUC1-ADTKD diagnosed via Probe extension Assay* | 44 copies normal 60 copies (1x dupC) |
| MUC1 p7 | 71 | kidney failure / 52yo | multigeneration family history of CKD; mother of p9 | diagnosed CKD 50yo with bland urinary sediment | MUC1-ADTKD diagnosed via Probe extension Assay* | 80 copies normal 83 x 60 (w. dupC) |
| MUC1 p8 | 41 | kidney failure / 41yo | multigeneration family history of CKD | presented with CKD at 27yo; bland urinary sediment | MUC1-ADTKD diagnosed via Probe extension Assay* | 45 copies normal 78 copies (1x dupC) |
| MUC1 p9 | 41 | CKD1 at 31yo | multigeneration family history of CKD; daughter of p7 | Diagnosed in context of family history; bland urinary sediment | MUC1-ADTKD diagnosed via Probe extension Assay* | 51 copies normal 83 copies (1x. dupC) |
| MUC1 p10 | 44 | kidney failure / 24yo | multigeneration family history of CKD | Presented in kidney failure; biopsy at presentation - severe tubulointerstitial disease | MUC1-ADTKD diagnosed via Probe extension Assay* | 40 copies normal 56 copies (1x dupC) |

CKD = Chronic Kidney Disease | * Method as per Kirby et al (2013)

#### SUPPLEMENTARY NOTE 3 – Genome assemblies for selected non-human vertebrates

To demonstrate that our Cornetto iterative genome assembly strategy is generalisable to diverse non-human species, we sequenced and assembled genomes for a selection of non-human vertebrates from different lineages. These were primarily selected for their high value in research and conservation. Together they serve to establish the suitability of Cornetto for assembling genomes of diverse architecture, such as genome and chromosome sizes, GC content, repeat content and heterozygosity, etc. Brief information on each species and details regarding sample collection, sequencing and assembly methods and outcomes are provided below.

#### Gould's petrel

Gould's petrel (*Pterodroma leucoptera*) is a threatened seabird with an extensive distribution across the Pacific Ocean, though breeding is limited to three small populations in Australia and New Caledonia. The main population of the Australian subspecies was successfully recovered from <250 pairs after intensive conservation measures including installing artificial nesting habitat, translocation and eradication of invasive species[14]. However, this population is again in decline and the reasons are unclear.

Petrel blood samples were collected from fledgling birds on Cabbage Tree Island, NSW on 9 March, 2024. Animal ethics permits were granted by the University of Tasmania Animal Ethics Committee (Project ID: 29495, approved 3 March, 2023) and blood was collected under scientific license granted under Part 2 of the Biodiversity Conservation Act 2016 by NSW Government Department of Climate Change, Energy, the Environment and Water (SL102834).

High–molecular weight (HMW) genomic DNA was extracted from a blood sample of one female individual and sequenced on a single PacBio SMRT cell. The resulting HiFi data was assembled to create a base genome assembly with metrics summarised below. The base assembly was augmented by running two ONT duplex flow cell in succession with Cornetto iterative sequencing (see **Methods**).

| Genome size | GC-content | Heterozygosity | Base asm data type | Base asm read length (N50) | Base asm read depth | N contigs in base asm (primary) |
|---|---|---|---|---|---|---|
| 1.4 Gbase | 43.8% | 0.7% | PacBio HiFi | 16300 | 55 | 411 |

#### Orange-bellied parrot

The orange-bellied parrot (*Neophema chrysogaster*) is a Critically Endangered migratory parrot species. Currently, the OBP is restricted to a single breeding site at Melaleuca, Tasmania, and its overwinter range is from east of the Murray River in South Australia and west of Port Phillip Bay in Melbourne[15]. The OBP is currently subject to intensive conservation management strategies, however, despite regular releases of juveniles from the captive population, the wild population is still small, numbering 81 individuals at the start of the 2023/24 breeding season[16]. Currently, the major threats to the species are ongoing habitat degradation, inbreeding, low genetic diversity and disease susceptibility[15,17,18].

Two captive adult male OBPs were euthanised on 20/08/2020 for medical reasons and immediately dissected for tissue collection. Tissues were collected for DNA sequencing (flash frozen at -80°C) under NSW Scientific Licence SL101204. High-molecular weight DNA was extracted from ~20mg liver tissue from one individual (SB# 2035) using the PacBio Nanobind kit standard Dounce homogenizer protocol. DNA was sequenced on two ONT flow cells (simplex data; LSK114). The resulting data was assembled to create a base genome assembly with metrics summarised below. The base assembly was then augmented by running one additional ONT flow cell with Cornetto iterative sequencing (see **Methods**).

| Genome size | GC-content | Heterozygosity | Base asm data type | Base asm read length (N50) | Base asm read depth | N contigs in base asm (primary) |
|---|---|---|---|---|---|---|
| 1.4 Gbase | 43.3% | 0.3% | ONT simplex | 32687 | 34 | 2084 |

Resolving the Major Histocompatibility Complex (*MHC*) region within the orange-bellied parrot genome is a critical priority. This locus is essential to gain an understanding and develop strategies to monitor the decline of immunogenetic diversity that threatens the survival of this species[19]. However, the MHC region is highly repetitive and therefore difficult to assemble. A recent genome assembly created with HiFi and HiC data was unable to resolve the MHC region[19]. To identify MHC genes in our updated Cornetto assembly, we used *blastn* with default parameters and publicly available MHC query sequences from Westerdahl et al. (2022)[20] and the orange-bellied parrot class I gene identified in Silver et al. (2025)[19]. BLAST results identified a single class I and alpha and beta class II gene in the assembly located on contig ptg000043l, within an 80kb genome region (between 180,000 and 260,000), with coordinates summarised below.

| Gene | Start | End | Strand |
|---|---|---|---|
| *MHCI* | 179703 | 182193 | - |
| *MHCII-A* | 254674 | 260679 | + |
| *MHCII-B* | 254179 | 250071 | - |

Therefore, in addition to significant improvements in the assembly contiguity and completeness, Cornetto successfully resolved the highly repetitive MHC region, which had eluded previous efforts.

#### Western saw-shelled turtle

The western saw-shelled turtle (*Myuchelys bellii*) is an endangered freshwater turtle (Family Chelidae). The species is restricted to high elevation streams in northern New South Wales and southern Queensland, Australia[21,22]. With restricted distribution and low abundance, the species is considered Endangered under the Australian national Environment Protection and Biodiversity Conservation (EPBC) Act 1999 and is the focus of a comprehensive conservation program to boost juvenile recruitment in wild populations[23]. Blood samples were collected from an adult male *M. bellii* sourced from George's Creek (-30° 8'11.08"S, 151° 23'5.01"E) for DNA sequencing on 18 December 2023 and stored at -80°C. Sample collection was conducted under authorization of the University of New England Animal Ethics Committee (ARA22-066) and under NSW Scientific Licence SL101876.

High-molecular weight DNA was extracted from blood using the PacBio Nanobind kit standard. DNA was sequenced on one ONT flow cell (simplex data; LSK114). The resulting data was assembled to create a base genome assembly with metrics summarised below. The base assembly was then augmented by running one additional ONT flow cell with Cornetto iterative sequencing (see **Methods**).

| Genome size | GC-content | Heterozygosity | Base asm data type | Base asm read length (N50) | Base asm read depth | N contigs in base asm (primary) |
|---|---|---|---|---|---|---|
| 2.0 Gbase | 43.4% | 0.4% | ONT simplex | 17720 | 57 | 228 |

#### Redstriped eartheater cichlid

The redstriped eartheater cichlid (*Geophagus surinamensis*) is a neotropical cichlid species exploited both as a food resource for local Amazonian populations and for the ornamental fish trade[24]. Blood samples were collected from an adult male specimen of this species in the Caripetuba River, Abaetetuba, Pará, Brazil (01°37’23.49”S, 048°55’33”W) on 19 May 2024. Sample collection was conducted under authorization from the Chico Mendes Institute for Biodiversity Conservation (ICMBio, Registration: 21078-13) and was approved by the Ethics Committee for the Use of Animals of the Federal University of Pará (Protocol: 8803211223-2023).

High–molecular weight (HMW) genomic DNA was extracted from a blood sample of one male individual and sequenced on a single PacBio SMRT cell. The resulting HiFi data was assembled to create a base genome assembly with metrics summarised below. The base assembly was augmented by running two ONT duplex flow cell in succession with Cornetto iterative sequencing (see **Methods**).

| Genome size | GC-content | Heterozygosity | Base asm data type | Base asm read length (N50) | Base asm read depth | N contigs in base asm (primary) |
|---|---|---|---|---|---|---|
| 1.0 Gbase | 41.3% | 0.6% | PacBio HiFi | 18843 | 78 | 142 |

#### SUPPLEMENTARY NOTE 4 - Extended Methods

### Example commands and configurations used for the analyses in the paper

##### Creating the base assembly

```
# for HiFi data:
hifiasm -t 48 --hg-size 3g -o asm-0 reads-0.fastq

# for ONT data (hifiasm version 0.22.0 or later is required:
hifiasm -t 48 --hg-size 3g --ont -o asm-0 reads-0.fastq
```

##### Getting the  boring bits for the first cornetto iteration

```
# if HiFi-based base assembly
minimap2 -t 24 --secondary=no --MD -ax map-hifi asm-0.fasta reads-0.fastq -o asm-0.sam

# if ONT-based base assembly
minimap2 -t 24 --secondary=no --MD -ax map-ont asm-0.fasta reads-0.fastq -o asm-0.sam


# sort and index
samtools sort -@ $24 asm-0.sam -o asm-0.bam
samtools index asm-0.bam

# get depths
samtools faidx asm-0.fasta
awk '{print $1"\t0\t"$2}' asm-0.fasta.fai | sort -k3,3nr > asm-0.chroms.bed
samtools depth -@ 24 -b asm-0.chroms.bed -aa asm-0.bam  | awk '{print $1"\t"$2-
1"\t"$2"\t"$3}' > asm-0.cov-total.bg
samtools depth -@ 24 -Q 20 -b asm-0.chroms.bed -aa asm-0.bam  | awk '{print $1"\t"$2-
1"\t"$2"\t"$3}' > asm-0.cov-mq20.bg


# run the create-cornetto script in the cornetto repository
scripts/create-cornetto.sh asm-0.fasta

# extra step for diploid assemblies
scripts/create-hapnetto.sh asm-0
```

##### Configuring readfish

```
# index the assembly
minimap2 -x map-ont -d asm-0.fasta.idx asm-0.fasta


# example of readfish .toml file

[caller_settings]
config_name = "dna_r10.4.1_e8.2_400bps_5khz_fast_prom"
host = "ipc:///tmp/.guppy"
port = 5555
align_ref = "/path/to/asm-0.fasta.idx"

[conditions]
reference = "/path/to/asm-0.fasta.idx"
[conditions.0]
name = "asm-0_dip.boringbits"
control = false
min_chunks = 0
max_chunks = 16
targets = "asm-0_dip.boringbits.txt"
single_on = "unblock"
multi_on = "unblock"
single_off = "stop_receiving"
multi_off = "stop_receiving"
no_seq = "proceed"
no_map = "proceed"
```

```
# launch readfish
readfish targets --device ${DEVICE_ID} --experiment-name asm-1 --toml asm-1.boringbits.toml
--port 9502 --cache-size 3000 --batch-size 3000 --channels 1 3000 --log-file my.log
```

**Basecalling ONT data for each cornetto iteration**

```
# For Duplex data
slow5-dorado duplex dna_r10.4.1_e8.2_400bps_sup@v4.2.0 reads.blow5  > reads.bam

# For Simplex data
slow5-dorado basecaller -x cuda:all dna_r10.4.1_e8.2_400bps_sup@v5.0.0 reads.blow5 --emit-
fastq --min-qscore 10  > reads_all.fastq

# filtering small reads (only for cornetto iterations using simplex data)
seqkit seq -m 30000 reads_all.fastq -o reads.fastq
```

**Assembling after each cornetto iteration**

```
# sequential reassembly for HiFi + duplex approach:
hifiasm -t 48 --hg-size 3g -o asm-n hifi_reads.fastq duplex_reads_batch1.fastq
duplex_reads_batch2.fastq duplex_reads_batchN.fastq

# sequential reassembly for ONT only approach:
hifiasm -t 48 --hg-size 3g --ont -o asm-n ont_reads_base.fastq ont_reads_batch1.fastq
ont_reads_batch2.fastq ont_reads_batchN.fastq

# re-generated boringbits target files for updated assemblies
scripts/recreate-cornetto.sh asm-n.fasta

# optional step for diploid assembly
scripts/create-hapnetto.sh asm-n
```

**Commands to run cornetto wrapper scripts for assembly evaluation (only for hg002)**

```
# index the Q100 assembly
# first download from:
#https://s3-us-west-2.amazonaws.com/human-
pangenomics/T2T/HG002/assemblies/hg002v1.0.1.fasta.gz
samtools faidx hg002v1.0.1.fasta.gz

# paternal haplotype with chrX
grep "PATERNAL\|chrEBV\|chrM\|chrX\|chrY" hg002v1.0.1.fasta.gz.fai | cut -f 1 >
paternal.txt
samtools faidx hg002v1.0.1.fasta.gz -r paternal.txt -o hg002v1.0.1_pat.fasta

# mapping and dotplot generation
scripts/minidotplot.sh hg002v1.0.1_pat.fasta asm.fasta

# get telomere counts
scripts/telostat.sh asm.fasta

# get per-chrosome statstics
scripts/asmstats.sh asm.fasta
```

**Command to get BUSCO scores**

```
# BUSCO scores
compleasm run -a asm.fasta -o asm.busco_out -t 8 -l ${LINEAGE} -L /path/to/mb_downloads
# LINEAGE: primates for human, actinopterygii_odb10 for cichlid, tetrapoda_odb10 for birds
and turtles
```

**Command to get QV, switch error and hamming error using yak (hg002 only)**

```
# create the yak index using the HG002 Q100 assembly, with a k-mer size of 21
yak count -k 21 -K1.5g -t 16 hg002v1.0.1.fasta.gz -o hg002v1.0.1.k21.yak
# create the yak index using the HG002 Q100 assembly, with a k-mer size of 31
yak count -k 31 -K1.5g -t 16 hg002v1.0.1.fasta.gz -o hg002v1.0.1.k31.yak

# QV (k=21) of primary ASM
```

```
yak qv hg002-k21.yak asm.fasta
# QV (k=31) of primary ASM
yak qv hg002-k31.yak asm.fasta

# QV of the combined haplotypes
cat asm.hap1.fasta asm.hap2.fasta > asm.combined.fasta
yak qv hg002-k21.yak asm.combined.fasta
yak qv hg002-k31.yak asm.combined.fasta

# Hamming and Switch errors
# download trio yak indexes from:
#https://s3-us-west-2.amazonaws.com/human-
pangenomics/index.html?prefix=submissions/6040D518-FE32-4CEB-B55C-504A05E4D662--
HG002_PARENTAL_YAKS/HG002_PARENTS_FULL
yak trioeval pat.HG003.yak mat.HG004.yak asm.combined.fasta -t 16
```

**Commands for NGA50, NGA90 and NGAx plots (hg002 only)**

```
# run quast
quast.py -t 48 -o hg002-cornetto-E_3.quast_out_align --large -r hg002v1.0.1.fasta
asm.combined.fasta
```

**Commands for structural variant calling and evaluation (hg002 only)**

```
# run dipcall on HG002 Q100 assembly to generate the truthset
dipcall dip chm13v2.0.fa hg002v1.0.1_pat.fasta hg002v1.0.1_mat.fasta  > dip.mak && make -j2
-f dip.mak

# run dipcall on our assembly
dipcall dip chm13v2.0.fa asm.hap1.fasta asm.hap2.fasta  > dip.mak && make -j2 -f dip.mak

# filtering for >50 base variants and indexing
bcftools norm -m-any dip.dip.vcf.gz > split.vcf
cat split.vcf | grep "^#" > structural_split.vcf
cat split.vcf | grep -v "^#" | awk '{if(length($4)>50 || length($5)>50) print $0}' >>
structural_split.vcf
bgzip structural_split.vcf  && tabix structural_split.vcf.gz

# evaluate
truvari bench -b hg002q100/structural_split.vcf.gz -c our_asm/structural_split.vcf.gz -f
chm13v2.0.fa -o output/
```

**Commands to remove non-human reads from saliva sample (before running hifiasm for the base assembly)**

```
# run centrifuge on FASTQ
centrifuge -p 48 -q -x p_compressed+h+v -U reads-0_all.fastq -S reads_classification.tsv --
report-file reads_report.tsv

# extract human reads
samtools faidx reads-0_all.fastq
awk '$3!=9606' reads_classification.tsv | cut -f 1 | sort -u > nonhuman_reads.txt
cut -f 1 reads-0_all.fai > all_reads.txt
grep -v -F -f nonhuman_reads.txt all_reads.txt > human_reads.txt
samtools fqidx -r human_reads.txt reads-0_all.fastq > reads-0.fastq
```

**Commands to get non-human contigs assembled from a saliva sample**

```
# identify nonhuman species with minimum 100 reads
sed 's/ /-/g' reads_report.tsv | sort -k5,5nr | awk '$2 != 9606' | awk '$5 >= 100' | cut -f
2 | sort -u | awk '$1 != "taxID"' > nonhuman_species_high_count.txt

# run centrifuge on FASTA
centrifuge -p 48 -f -x p_compressed+h+v -U asm-all-0.fasta -S contig_classification.tsv --
report-file contig_report.tsv
```

**Commands to measure compute resources**

```
# Elapsed (wall clock) time, Max RAM (resident set size)
/usr/bin/time -v hifiasm …
```

```
# disk usage
du -h <file>
```

**Commands for hg002-Cornetto-4 assemblies with and without HiC data**

```
hg002-Cornetto-4 (partially phased asm) contigs
hifiasm --ont -t 48 --telo-m CCCTAA --hg-size 3g -o asm D_0_PGXXSX240470.fastq E_1_QGXXXX250049.fastq E_2_QGXXXX250062.fastq E_3_QGXXXX250063.fastq

hg002-Cornetto-4 (partially phased asm) scaffolds
hifiasm --ont -t 48 --telo-m CCCTAA --dual-scaf --hg-size 3g -o asm D_0_PGXXSX240470.fastq E_1_QGXXXX250049.fastq E_2_QGXXXX250062.fastq E_3_QGXXXX250063.fastq

hg002-Cornetto-4-HiC (fully phased asm) contigs
R1=HG002.HiC_2_NovaSeq_rep1_run2_S1_L001_R1_001.fastq.gz
R2=HG002.HiC_2_NovaSeq_rep1_run2_S1_L001_R2_001.fastq.gz
hifiasm --ont -t 48 --telo-m CCCTAA --hg-size 3g -o asm --h1 ${R1} --h2 ${R2} D_0_PGXXSX240470.fastq E_1_QGXXXX250049.fastq E_2_QGXXXX250062.fastq E_3_QGXXXX250063.fastq

hg002-Cornetto-4-HiC (fully phased asm) scaffolds
hifiasm --ont -t 48 --telo-m CCCTAA --dual-scaf --hg-size 3g -o asm --h1 ${R1} --h2 ${R2} D_0_PGXXSX240470.fastq E_1_QGXXXX250049.fastq E_2_QGXXXX250062.fastq E_3_QGXXXX250063.fastq
```

#### Software versions used for analyses in this paper

- cornetto: 0.1.0-alpha
- hifiasm: 0.19.8 for HiFi data
- hifiasm: 0.22.0 for ONT data; 0.25.0 for hg002-Cornetto-4
- slow5-dorado: 0.3.4 for duplex, 0.8.3 for simplex
- minimap2: 2.24
- compleasm: 0.2.6
- yak: 0.1-r69-dirty
- centrifuge: 1.0.4
- seqkit: 2.3.0
- quast: 5.2.0
- dipcall: 0.3
- bcftools: 1.16
- truvari: 5.3.0
